# Training a spiking neuronal network model of visual-motor cortex to play a virtual racket-ball game using reinforcement learning

**DOI:** 10.1101/2021.07.29.454361

**Authors:** Haroon Anwar, Simon Caby, Salvador Dura-Bernal, David D’Onofrio, Daniel Hasegan, Matt Deible, Sara Grunblatt, George L Chadderdon, Cliff C Kerr, Peter Lakatos, William W Lytton, Hananel Hazan, Samuel A Neymotin

## Abstract

Recent models of spiking neuronal networks have been trained to perform behaviors in static environments using a variety of learning rules, with varying degrees of biological realism. Most of these models have not been tested in dynamic visual environments where models must make predictions on future states and adjust their behavior accordingly. The models using these learning rules are often treated as black boxes, with little analysis on circuit architectures and learning mechanisms supporting optimal performance.

Here we developed visual/motor spiking neuronal network models and trained them to play a virtual racket-ball game using several reinforcement learning algorithms inspired by the dopaminergic reward system. We systematically investigated how different architectures and circuit-motifs (feed-forward, recurrent, feedback) contributed to learning and performance. We also developed a new biologically-inspired learning rule that significantly enhanced performance, while reducing training time.

Our models included visual areas encoding game inputs and relaying the information to motor areas, which used this information to learn to move the racket to hit the ball. Neurons in the early visual area relayed information encoding object location and motion direction across the network. Neuronal association areas encoded spatial relationships between objects in the visual scene. Motor populations received inputs from visual and association areas representing the dorsal pathway. Two populations of motor neurons generated commands to move the racket up or down. Model-generated actions updated the environment and triggered reward or punishment signals that adjusted synaptic weights so that the models could learn which actions led to reward.

Here we demonstrate that our biologically-plausible learning rules were effective in training spiking neuronal network models to solve problems in dynamic environments. We used our models to dissect the circuit architectures and learning rules most effective for learning. Our model shows that learning mechanisms involving different neural circuits produce similar performance in sensory-motor tasks. In biological networks, all learning mechanisms may complement one another, accelerating the learning capabilities of animals. Furthermore, this also highlights the resilience and redundancy in biological systems.

## Introduction

A variety of Deep Learning (DL) artificial neural network (ANN) models have been developed and trained to effectively learn complex sensory-motor behaviors [1–8]. DL models, which are primarily designed with engineering goals in mind, often lack biological details, and therefore do not shed light on the circuit mechanisms of behavior in real animals [9].

Biophysically detailed neuronal network models of the sensory-motor cortex can be used to dissect the mechanisms of learning behaviors in vivo [10], however in the past the focus has been on developing models of cortical circuits that reproduce electrophysiological activity patterns [11] rather than on understanding the origins of sensorimotor behavior [12]. Several spiking neuronal networks (SNNs) with moderate circuit complexity have been developed to learn behaviors in static sensory environments [13–17]. In this work, we aim to shed light on the dynamics, decision-making, and learned behavior of a visual-motor circuit in a dynamic environment. We develop several SNNs each including multiple visual and motor areas that learn to interact with the environment using biologically inspired reinforcement learning (RL) mechanisms.

The success of ANNs can be credited to the backpropagation and gradient descent methods that successfully tune the connection weights between neurons [18]. From a biological perspective, the ideas behind the backpropagation and gradient descent methods are very appealing but many properties and requirements that it relies on to tune synaptic connections are not present in the nervous system [19]. Nevertheless, the success of ANNs with backpropagation and gradient descent has led to achieving superhuman capabilities in learning goals and learning to operate in an interactive environment [5]. One of the techniques that have been used to train ANNs interacting with an environment is reinforcement learning (RL), where the network learns a behavior by maximizing a reward signal from the environment. Our goal is not to compete with the success of ANNs (although the success of the model is important), but rather to improve understanding of the intricate networks of the visual and motor systems that learn using biologically realistic time-dependent and reward modulated learning rules. Using the biologically inspired RL rule we can not only show that our models perform well but also that their neuronal activity is directly comparable to recordings of biological networks.

Cortical neural circuits contain very complex connectivity patterns [20–22]. Sensory areas are connected with one another and to motor and other higher processing areas using multiple pathways [23–29]. In addition to feedforward connections, feedback and recurrent connections are hallmarks of biological neural circuits. However, it remains unclear what role each of those connections serve in neural computations, in multimodal integration of sensorimotor information, and in generating motor behavior. Using SNN models with feedforward, recurrent, and feedback connections in this work, we investigate learning capacity of these models with different architectures and connectivity patterns.

In sensory-motor tasks, rewards and punishments are typically sparsely delivered at the end of each trial, where each trial consists of multiple actions in a dynamically changing environment [30, 31]. The brain utilizes environmental cues in dynamically changing environments to make associations with the actions that eventually result in a reward over repeated trials [32]. Regardless of the temporal delays between the executed actions and rewards, the brain is capable of assigning credit to intermediate actions during a trial. Several theoretical solutions have been proposed to solve this distal credit assignment problem in both ANN and SNN models [33–36]. Reinforcement learning in ANNs has made heavy use of value functions to assign intermediate credit in sparse reward paradigms, but this methodology remains impractical in biological SNNs [34,37–39]. In this work, we use a spike-timing-dependent plasticity (STDP) rule to establish association between pre- and postsynaptic neurons, and modulate the STDP weight changes by reward/punishment (*critic signal*) delivered after an action. Besides these types of temporally *sparse*, delayed rewards, we also test another reward paradigm utilizing *intermediate* rewards, which were previously used with an SNN model of sensorimotor cortex trained to move simulated and robotic arms towards targets [14, 40]. However, instead of broadcasting *critic* signals to all premotor and motor neuron pairs (*non-targeted paradigm*), we provide *intermediate* rewards/punishments only to the neuronal populations associated with the executed actions (*targeted paradigm*).

In this work, we first construct a feedforward SNN model of visual and motor areas and train it to play a racket-ball game using STDP based RL (STDP-RL) mechanisms with intermediate rewards/punishment in both *targeted* and *non-targeted* RL paradigms. We then extend our feedforward model to include feedback and recurrent connections, as well as allowing RL based learning within premotor and motor areas. Like the feedforward model, we also test our recurrent model’s ability to learn under both *targeted* and *non-targeted* RL paradigms using both *intermediate* and *sparse* rewards. Comparing performance of our models using both feedforward and recurrent architectures with different RL paradigms, we show the capability of SNN models in learning complex visual-motor behaviors, which were previously demonstrated only using ANNs. Furthermore these models allow us to access the spiking activity of neurons across different modeled areas that can be directly matched to physiological data. Once more anatomical and physiological details about neural circuits and RL modalities are included in our models, these models can be used together with imaging modalities to dissect the mechanisms of psychiatric disorders associated with deficits in sensory-motor behaviors [41–43].

## Materials and Methods

### “Racket-ball” game

We designed a “Racket-ball” game to use for training our visual-motor cortex model to play. Many features of the game were designed to resemble those of the Atari games (especially Pong except that there was no opponent; (see Image frames in **Figure 1A and 7A**) provided by OpenAI’s gym platform (https://gym.openai.com) [44]. In a court (160 pixels x 160 pixels), the racket (4 pixels wide and 16 pixels high) was controlled by external motor commands to move vertically up and down at a fixed horizontal position (140th pixel). At every new serve, the ball’s (4 pixels x 4 pixels) position was reset to the extreme left side of the court with a randomly selected vertical position (possible vertical starting locations of the ball: 40,60,80,100,120 pixels). However, the vertical position of the ball and the racket at the first serve of each episode could be specified externally. The motion direction and speed of the ball at each serve was randomly initiated by choosing the displacement in horizontal and vertical direction from (dx,dy) = {(1,1),(1,2),(1,3),(2,1),(2,2),(2,3),(3,1),(3,2),(3,3)}. When the ball hit the upper or lower edge of the court, it bounced back in the vertical direction (-dy) without any change in the speed or the horizontal direction (dx). Every time when the ball was hit by the racket, it bounced back in a different direction depending on the contact point of the ball with the racket. When the ball was hit by the racket’s lower or upper edge, the ball bounced back with double speed in the new randomly selected vertical direction (dy). Similar to the ball bouncing after hitting the left edge of the court, when the ball was hit by the racket’s center, the ball bounced back only with the new randomly selected vertical direction. When the ball was hit by the racket, a point was awarded (+1) and when the ball was missed by the racket, a point was deducted (-1) from the total score.

**Figure 1.**
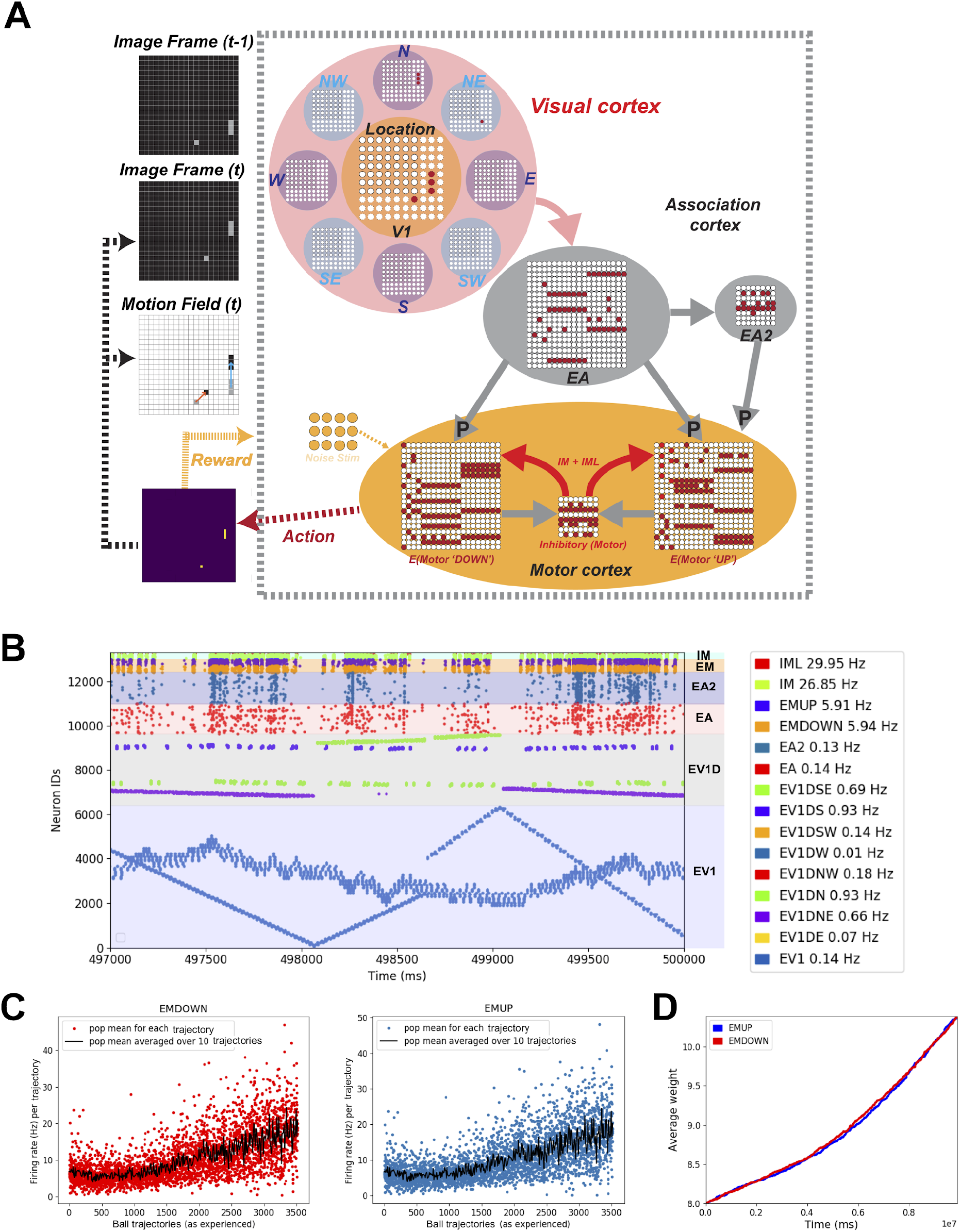
Constructing a feedforward model of visual-motor cortex that learns to play the racket-ball game. **A)** Schematic of the closed-loop feedforward visual/motor circuit model interfaced with the racket-ball game. Visual areas receive inputs from the pixelated image frames of the racket-ball game, downstream activating association and motor areas. An action is generated after comparing firing rates of EMDOWN and EMUP excitatory motor populations over an interval. Each action delivers a reward to the model driving STDP-RL learning rules. **B)** Raster plot shows the spiking activity of different populations of neurons during a training episode (vertical axis is neuron identity and horizontal axis is time; each dot represents a single action potential from an individual neuron). Patterned activation of early visual neurons (i.e. diagonal lines in raster plots) indicate for example, the ball traversing the court from side to side. These patterns are visible because the early visual neurons were arranged topographically with increasing neuron number. **C)** Firing rates of excitatory motor neuron populations EMUP and EMDOWN in the feedforward model increase over the course of training. The firing rates were binned for ball trajectories (beginning when the ball is at the extreme left side of the court and ends when the ball hits or misses the racket on the right side of the court). **D)** The average weight change of synaptic input onto EMUP and EMDOWN sampled over 20 training episodes tends to increase with learning, indicating the network tends to produce rewarding behavior.

### Intermediate Reward paradigm

In standard game environments, sparse rewards and punishments are awarded based on scoring or losing a point. But while learning how to play a game, all the correct moves/actions during the time available to respond according to the situation contribute towards the end result i.e. whether a player scores a point or not. In a “Racket-ball” game such moves could be making a proper serve, estimating the projection of the ball after bouncing back towards the player, and taking a proper action/move toward the estimated contact position. Also if the player makes an incorrect move, they could compensate for the mistake and move in the correct direction in the next step. All these actions or movements during training eventually lead to the player becoming an expert over repeated episodes or matches. Based on these intuitive learnable cues, in addition to the standard reward and punishment, we proposed using intermediate rewards i.e. award a small reward (+0.1) or smaller punishment (-0.01) at each action the player takes based on whether that action contributed in moving the racket towards the projected position of the ball to be hit or not. The schematic of the intermediate reward paradigm is shown in **Figure 2B**. When the ball moved towards the racket, we used the direction of the motion of the ball to predict the potential position of the ball where the racket could hit it. Using this projected position, when the racket moved towards the target position, we awarded a small reward (+0.1) for making the correct move and when the racket moved away from the target position, we awarded a smaller punishment (-0.01) for making an incorrect move.

**Figure 2:**
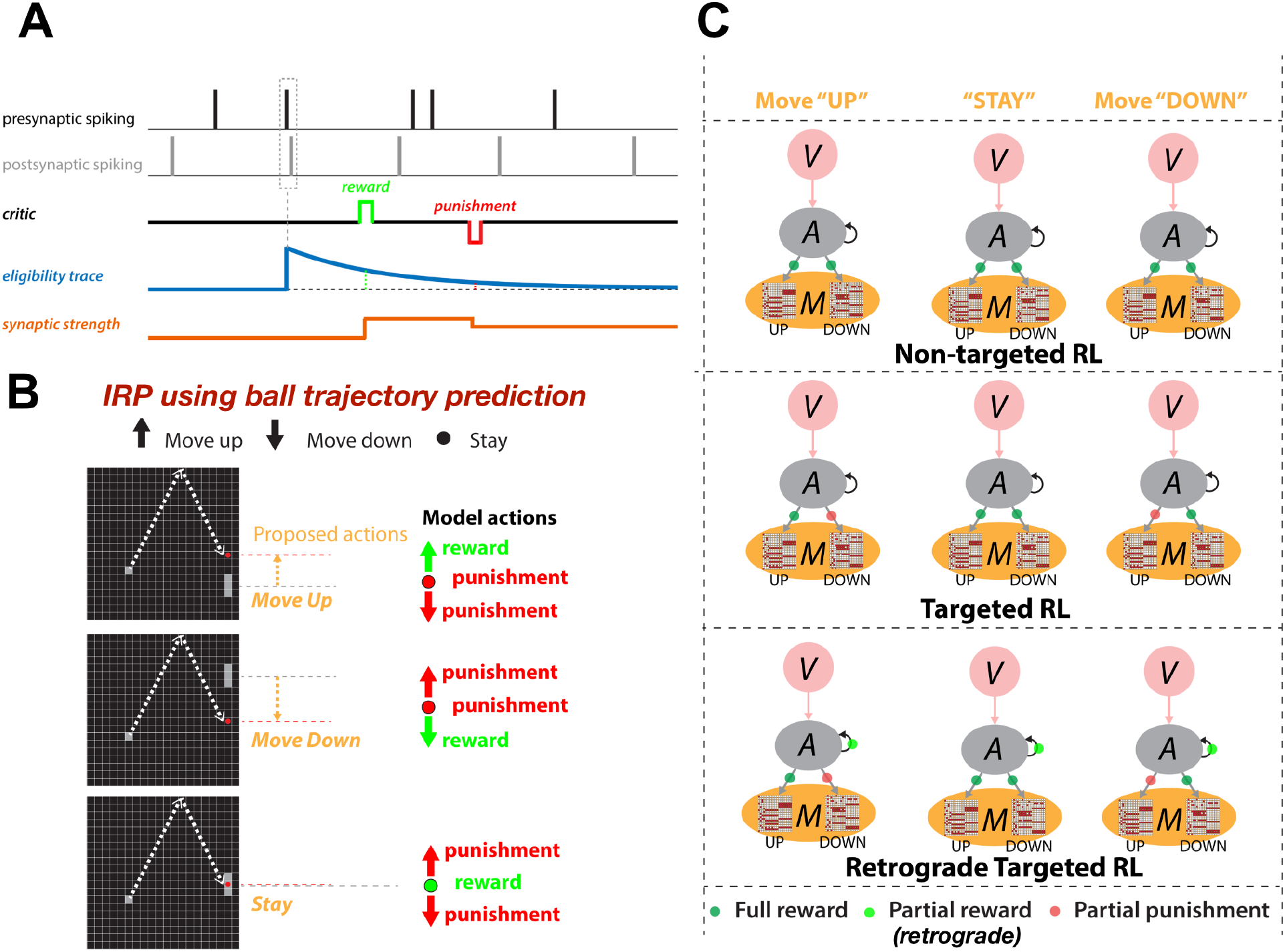
Spike-timing dependent reinforcement learning framework: **A)** An exponentially-decaying synaptic eligibility trace (ET) is triggered after postsynaptic neuron firing within a short time window after presynaptic neuron firing. If a reward or punishment signal is delivered while ET>0, synaptic weight is potentiated or depressed proportional to ET. **B)** IRP delivers rewards to the model for each action it takes based on whether the action moved the racket towards the projected location of the ball for a hit or away. **C)** Three different RL versions used in this study (*V* visual; *A* Association; *M* Motor areas): ***non-targeted RL***, all motor neurons receive ET; ***targeted RL***, motor neurons which contributed to the action receive ET and motor neurons in population for the other directions receive negative ET; ***retrograde targeted RL*** as in ***t****argeted RL* but middle/hidden layer synaptic connections also receive ET, with ET amplitude reduction based on number of back-tracked connections.

It should be noted that no explicit loss function is used in our SNN in contrast to traditional ANNs. Rather, it is expected that the total rewards will be maximized with the defined learning framework without the internal drive to minimize a loss function.

### Reinforcement learning paradigms

We used the existing spike-time-dependent reinforcement learning (STDP-RL) mechanism in this work, which was developed based on the distal reward learning paradigm proposed by Izhikevich [33], with variations used in neuronal network models [13,14,45,46]. Our STDP-RL used a spike-time-dependent plasticity mechanism together with reward or punishment signal for potentiation or depression of the targeted synapses. An exponentially decaying eligibility trace was included to assign temporally distal credits to the relevant synaptic connections. The term “eligibility trace” is used to define a time window during which an eligible synaptic connection between a pair of neurons can undergo strengthening or weakening.

Historically this term has been used in three-factor learning rules, where the eligibility trace usually allows a synapse to undergo a plastic change far in time from when the neuromodulator becomes available. Experimental evidence for eligibility traces in striatum [47], cortex [48], and hippocampus [49–51] shows distinct temporal dynamics ranging from 1 to 5 seconds across different areas. On the molecular level, the striatal three-factor plasticity depends on NMDA, CaMKII, protein synthesis and dopamine D1 receptors [47, 52]. CaMKII increases were found to be localized in dendritic spines and to have roughly the same time course as the critical window for phasic dopamine. This suggests CaMKII could be involved in the “synaptic flag” triggered by the STDP-like induction protocol. Protein kinase A (PKA) was found to have a nonspecific cell-wide distribution suggesting an interpretation of PKA as a dopamine-triggered third factor [47]. The STDP-RL mechanism used in this work is depicted in **Figure 2A**: when a postsynaptic spike occurred within a few milliseconds of the presynaptic spike, the synaptic connection between this pair of neurons became eligible for STDP-RL and was tagged with an exponentially decaying eligibility trace. Later, when a reward or a punishment was delivered before the eligibility trace decayed to zero, the weight of the tagged synaptic connection was increased or decreased, depending on the ‘critic’ value and sign i.e. increase for reward or decrease for punishment. The change in synaptic strength was proportional to the eligibility trace value at the time of the critic’s delivery (see **Figure 2A**).

Traditionally, when using STDP-RL for learning behavior, all plastic synaptic connections in the neuronal network model are treated equally considering that the underlying causality between pre and postsynaptic neurons and the associated action and critic will automatically choose only relevant synaptic connections for reinforcement. We used this standard STDP-RL (see *non-targeted RL* in **Figure 2C**) approach in this study, but also proposed several variations to this standard approach, which required the presence of distinct populations of neurons controlling distinct behaviors. In the first variation (see *targeted RL* in **Figure 2C**), we delivered full reward or punishment to the neuronal population that generated the action and additionally, we delivered opposite and partial ‘critic’ value to the non-action associated neuronal population. This ensured that the learning happened only in the part of the circuit which generated the action. In the second variant (see *retrograde targeted RL* in **Figure 2C**), we further extended the partial ‘*critic*’ value delivery to the neuronal populations one synapse away from those directly generating motor action.

The parameters of the STDP-RL were adjusted to incorporate temporally well-separated motor actions, visual scenes and associated rewards. For intermediate scenarios and associated rewards, shorter time constants were sufficient to allow learning those intermediate level performances.

### Excitatory and Inhibitory neurons used in the model

All modeled neurons were point neurons, and did not include different soma or dendrite compartments. However, in order to allow more dynamic complexity in our modeled neurons, we included state variables and synaptic inputs that represented activation at soma or distal dendrites. This was done by using longer synaptic delays and longer synaptic time constants from pre- and post-synaptic locations that represented dendrite vs soma synapses. These details are fully described in our previous papers [53]. Individual neurons were modeled as event-driven, rule-based dynamical units with many of the key features found in real neurons, including adaptation, bursting, depolarization blockade, and voltage-sensitive NMDA conductance [54–57]. Event-driven processing provides a faster alternative to network integration: a presynaptic spike is an event that arrives after a delay at a postsynaptic neuron; this arrival is then a subsequent event that triggers further processing in the postsynaptic neurons. Neurons were parameterized as excitatory (E), fast-spiking inhibitory (I), and low voltage activated inhibitory (IL; Table 1). Each neuron had a membrane voltage state variable (V_m_), with a baseline value determined by a resting membrane potential parameter (V_rest_). After synaptic input events, if V_m_ crossed spiking threshold (V_thresh_), the cell would fire an action potential and enter an absolute refractory period, lasting τ_AR_ ms. After an action potential, an after-hyperpolarization voltage state variable (V_AHP_) was increased by a fixed amount ΔV_AHP_ and then V_AHP_ was subtracted from V_m_. Then V_AHP_ decayed exponentially (with time constant τ_AHP_) to 0. To simulate depolarization blockade, a neuron could not fire if V_m_ surpassed the blockade voltage (V_block_). Relative refractory period was simulated after an action potential by increasing the firing threshold V_thresh_ by W_RR_(V_block_-V_thresh_), where W_RR_ was a unitless weight parameter. V_thresh_ then decayed exponentially to its baseline value with a time constant τ_RR_.

**Table 1:** Parameters of the neuron model for each type.

| Cell type | $V_{rest}$ (mV) | $V_{thresh}$ (mV) | $V_{block}$ (mV) | $\tau_{AR}$ (ms) | $W_{RR}$ | $\tau_{RR}$ (ms) | $\Delta V_{AHP}$ (mV) | $\tau_{AHP}$ (ms) |
| --- | --- | --- | --- | --- | --- | --- | --- | --- |
| Excitatory (E) | -65 | -40 | -25 | 5 | 0.75 | 8 | 1 | 400 |
| Inhibitory (I) | -63 | -40 | -10 | 2.5 | 0.25 | 1.5 | 0.5 | 50 |
| Low-threshold Inhibitory (IL) | -65 | -47 | -10 | 2.5 | 0.25 | 1.5 | 0.5 | 50 |

$V_{rest}$ =resting membrane potential; $V_{thresh}$ =spiking threshold, $V_{block}$ =depolarization blockade voltage, $\tau_{AR}$ =absolute refractory time constant, $W_{RR}$ =relative refractory weight, $\tau_{RR}$ =relative refractory time constant, $\Delta V_{AHP}$ =after-hyperpolarization increment in voltage, $\tau_{AHP}$ =after-hyperpolarization time constant.

### Synaptic mechanisms

In addition to the intrinsic membrane voltage state variable, each cell had four additional voltage state variables V_syn_ corresponding to the synaptic inputs. These represent AMPA (AM2), NMDA (NM2), and somatic and dendritic GABAA (GA and GA2) synapses. At the times of input events, synaptic weights were updated by step-wise changes in V_syn_, which were then added to the cell’s overall membrane voltage V_m_. To allow for dependence on V_m_, synaptic inputs changed V_syn_ by dV=W_syn_(1-V_m_/E_syn_), where W_syn_ is the synaptic weight and E_syn_ is the reversal potential relative to V_rest_. The following values were used for the reversal potential E_syn_: AMPA, 65 mV; NMDA, 90 mV; and GABAA, –15 mV. After synaptic input events, the synapse voltages V_syn_ decayed exponentially toward 0 with time constants τ_syn_. The following values were used for τ_syn_: AMPA, 10 ms; NMDA, 300 ms; somatic GABAA, 10 ms; and dendritic GABAA, 20 ms. The delays between inputs to dendritic synapses (AMPA, NMDA, dendritic GABAA) and their effects on somatic voltage were selected from a uniform distribution ranging between 3– 5 ms, while the delays for somatic synapses (somatic GABAA) were selected from a uniform distribution ranging from 1.8–2.2 ms. Synaptic weights were fixed between a given set of populations except for those involved in learning (see RL “on” or “off” in Tables 2 and 3).

**Table 2:** Area interconnection probabilities and initial weights for the feedforward model.

| Presynaptic type | Postsynaptic type | Synapse type | Connection Convergence | Synaptic Weight | RL plasticity |
| --- | --- | --- | --- | --- | --- |
| EV1 | EA | AM2 | 128 | 6 | Off |
|  |  | NM2 |  | 0.1 | Off |
| EV1D | EA | AM2 | 8 | 6 | Off |
|  |  | NM2 |  | 0.1 | Off |
| EA | EA2 | AM2 | 30 | 12.5 | Off |
|  |  | NM2 |  | 0.15 | Off |
|  | EM | AM2 | 30 | 12 | On |
|  |  | NM2 |  | 0.15 | Off |
| EA2 | EM | AM2 | 30 | 12 | On |
| EM | IM | AM2 | 20 | 10 | Off |
|  |  | NM2 |  | 0.0195 | Off |
|  | IML | AM2 | 20 | 5 | Off |
|  |  | NM2 |  | 0.098 | Off |
| IM | EM | GA | 22 | 9 | Off |
| IML | EM | GA2 | 8 | 2.5 | Off |

**Table 3:** Area interconnection probabilities and initial weights for the recurrent model.

| Presynaptic type | Postsynaptic type | Synapse type | Connection Convergence | Synaptic Weight | RL plasticity |
| --- | --- | --- | --- | --- | --- |
| EV1 | EA | AM2 | 640 | 12.5 | Off |
|  |  | NM2 |  | 1 | Off |
|  | EA2 | AM2 | 30 | 0.25 | Off |
|  | EM | AM2 | 30 | 0.25 | Off |
| EA | EA | AM2 | 30 | 0.05 | On |
|  |  | NM2 |  | 0.005 | Off |
|  | EA2 | AM2 | 100 | 0.5 | On |
|  |  | NM2 |  | 0.01 | Off |
|  | EM | AM2 | 100 | 0.5 | On |
|  |  | NM2 |  | 0.01 | Off |
|  | IA | AM2 | 93 | 1.95 | Off |
|  |  | NM2 |  | 0.98 | Off |
|  | IAL | AM2 | 110 | 0.98 | Off |
|  |  | NM2 |  | 0.098 | Off |
| EA2 | EA | AM2 | 3 | 0.05 | On |
|  |  | NM2 |  | 0.005 | Off |
|  | EA2 | AM2 | 60 | 0.5 | On |
|  |  | NM2 |  | 0.01 | Off |
|  | EM | AM2 | 60 | 0.5 | On |
|  |  | NM2 |  | 0.01 | Off |
|  | IA2 | AM2 | 93 | 1.95 | Off |
|  |  | NM2 |  | 0.0195 | Off |
|  | IA2L | AM2 | 110 | 0.98 | Off |
|  |  | NM2 |  | 0.098 | Off |
| EM | EM | AM2 | 60 | 0.5 | On |
|  |  | NM2 |  | 0.01 | Off |
|  | EA | AM2 | 3 | 0.05 | On |
|  |  | NM2 |  | 0.005 | Off |
|  | EA2 | AM2 | 3 | 0.05 | On |
|  |  | NM2 |  | 0.005 | Off |
|  | IM | AM2 | 93 | 1.95 | Off |
|  |  | NM2 |  | 0.0195 | Off |
|  | IML | AM2 | 110 | 0.98 | Off |
|  |  | NM2 |  | 0.098 | Off |
| IM | EM | GA | 22 | 9 | Off |
|  | IM | GA | 31 | 4.5 | Off |
|  | IML | GA | 2 | 4.5 | Off |
| IA | EA | GA | 22 | 9 | Off |
|  | IA | GA | 31 | 4.5 | Off |
|  | IAL | GA | 17 | 4.5 | Off |
| IAL | EA | GA2 | 8 | 2.5 | Off |
|  | IA | GA2 | 12 | 2.5 | Off |
|  | IAL | GA2 | 2 | 4.5 | Off |
| IA2 | EA2 | GA | 22 | 9 | Off |
|  | IA2 | GA | 31 | 4.5 | Off |
|  | IA2L | GA | 17 | 4.5 | Off |
| IA2L | EA2 | GA2 | 8 | 2.5 | Off |
| IA2L | IA2 | GA2 | 12 | 2.25 | Off |
| IA2L | IA2L | GA2 | 2 | 4.5 | Off |

### Constructing spiking visual-motor cortex models for reward based learning

We built several versions of the spiking network models of the visual-motor cortex which can be mainly grouped into *feedforward* and *recurrent* models. Both models used excitatory (E) and inhibitory (I and IL) neurons with same excitability properties and synaptic dynamics. As suggested by the name, the main difference between the two types of models is the connectivity patterns and the synaptic weights in the network. Based on the distinctive features of the model, we denoted the first model as “feedforward model” and the other model as “recurrent model”.

#### The feedforward model

In the feedforward model (**Figure 1A**), we included several populations of neurons in the visual cortex model, one for encoding spatial location and the other 8 populations for encoding motion direction of the objects. Similar to the recurrent model, we also included two layers of object-associating neurons representing the association cortex and a single layer with two distinct populations of motor neurons representing the motor cortex. In the visual cortex model, we included 6400 location encoding E neurons (EV1) and 8 populations of 400 direction specific neurons (EV1D) each. We increased the association space as compared to the recurrent model by including 1400 E neurons in each layer (EA and EA2) of association cortex model. For the motor cortex model (M), we used two populations of 300 E neurons each (EMUP and EMDOWN) representing the motor areas generating “UP” and “DOWN” motor commands. We included inhibitory neurons (206 IM and 94 IML) only in the motor cortex.

Each EV1 neuron received a single input from a spike-generating Poisson process driven by individual pixels in the input image at 20Hz, with a delay of 1.8 to 2.2 ms (uniform distribution). Similarly, each EV1D neuron received a single input from a spike-generating Poisson process driven by the direction of object motion at individual pixels in the input image. Each EA neuron received excitatory inputs from 128 randomly selected EV1 and 8 randomly selected EV1D neurons from each of 8 direction selective populations (responsive to movements West, Northwest, North, Northeast, East, Southeast, South, Southwest). Each EA2 neuron received excitatory inputs only from 30 EA neurons. Each EM neuron received excitatory inputs from 300 EA and 300 EA2 neurons, and inhibitory inputs from 22 IM and 8 IML neurons. Each IM neuron received excitatory input from 20 neurons of each EM subpopulation. Each IML neuron received excitatory input from 20 neurons of each EM subpopulation. All neurons received excitation through AMPA (AM2) and NMDA(NM2) synapses. Only motor neurons received inhibition from IM and IML using GABAA synaptic mechanisms (GA and GA2). See Table 2 for all the initial synaptic connection weights.

#### The recurrent model

In the recurrent model (**Figure 7A**), we included a single layer of neurons encoding spatial location in the visual cortex, two layers of object-associating neurons representing association cortex and a single layer with two distinct populations of motor neurons representing motor cortex. We included 6400 E neurons in the visual cortex model (V1), 600 E neurons in the first layer of association cortex model (A) and 300 E neurons in the second layer of association cortex model (A2). For the motor cortex model (M), we used two populations of 300 E neurons each (EMUP and EMDOWN) representing the motor areas generating “UP” and “DOWN” motor commands. We used two types of inhibitory neurons in the network, 206 I and 94 IL neurons in A, 103 I and 47 IL neurons in A2, and 206 I and 94 IL neurons in M. We did not include any inhibitory neurons in V1.

Each E neuron in V1 received a single input from a spike-generating poisson process driven by individual pixels in the input image at 35 Hz, with a delay of 1.8 to 2.2 ms. Each EA neuron received excitatory inputs from 640 randomly selected EV1 neurons (feedforward E), 30 randomly selected EA neurons (recurrent E), 3 randomly selected EA2 neurons (feedback E) and 3 randomly selected EMUP and EMDOWN neurons each (feedback E), and inhibitory inputs from 22 IA and 8 IAL neurons. Each IA neuron received excitatory inputs from randomly selected 93 EA neurons and inhibitory inputs from randomly selected 31 IA (recurrent I) and 12 IAL neurons. Each IAL neuron received excitatory inputs from randomly selected 110 EA neurons and inhibitory inputs from randomly selected 17 IA and 2 IAL neurons. Each EA2 neuron received excitatory inputs from 30 EV1, 100 EA, 60 EA2, 3 EMUP and 3 EMDOWN neurons, and inhibitory inputs from 22 IA2 and 8 IA2L neurons. Each IA2 neuron received excitatory inputs from 93 EA2 neurons and inhibitory inputs from 31 IA2 and 12 IA2L neurons. Each IA2L neuron received excitatory inputs from 110 EA2 neurons and inhibition from 17 IA2 and 2 IA2L neurons. Each EM neuron received excitatory inputs from 30 EV1, 100 EA, 60 EA2 and 60 EM neurons, and inhibitory inputs from 22 IM and 8 IML neurons. Additionally, each EM subpopulation receive reciprocal inhibition from the other EM subpopulation with probability of 0.125. Each IM neuron received excitatory input from 93 neurons of each EM subpopulation and inhibitory inputs from 31 IM and 12 IML neurons. Each IML neuron received excitatory input from 110 neurons of each EM subpopulation and inhibitory inputs from 17 IM and 2 IML neurons. Each excitatory synaptic input was implemented using both AMPA and NMDA synapses. See Table 3 for all initial synaptic connection weights.

The number of interneurons of a specific type, for example IM and IML, were based on our previous models of neocortex where a proper balance between the number of fast spiking IM and low-threshold spiking IML are required to prevent either too much or too little inhibition in the circuit [14]. Since the IML neurons more readily fire, their number is lower. There is also evidence in neocortical circuitry of these distinctions in the number of interneurons of different types. For example, fast-spiking PV interneurons (IM in our model) which target soma have higher density in the neocortex compared to the SOM dendrite-targeting interneurons, which in our model are simulated using the IML population.

The number of inputs from specific populations were chosen to replicate the somewhat sparse connection densities, which are seen in the neocortex, and are based on our previous models, as cited. However, we note that the exact values are not intended to represent values that would be measured from specific circuit mapping studies, which would be difficult to constrain given that our number of populations are vastly simplified compared to the dozens of neuronal types seen in neocortex. Instead, we aimed to use rough values observed from circuit mapping studies.

### Generating motor commands

We built a modular model of motor areas specific to the “Racket-ball” game, which required 3 motor commands (Move Up, Move Down, Stay). To encode these commands, we used 2 motor areas, associated with “Move UP” or “Move Down”. The output motor command was generated from the motor area based on a *winner takes all* rule, meaning motor commands were determined by the maximum population-firing rates across motor areas e.g. if the population-firing rate of motor area representing motor command “Move Up” was larger than the population-firing rate of motor area representing motor command “Move Down”, then “Move Up” command would be generated. If the population-firing rate of motor areas representing both motor commands is the same, the third motor command “Stay” was generated. Each action in the feedforward model was produced every 20 ms interval, whereas the recurrent model used 50 ms intervals.

### Interfacing visual-motor cortical model with the “Racket-ball” game

We interfaced our visual-motor cortex model with the custom built game environment “Racket-ball”, allowing the model to sense and act on visual information from the game. At each game-step (20 ms for the feedforward model and 50 ms for the recurrent model), the model read screen pixels from an image frame, processed information and generated a motor command. In return this produced an intermediate reward (0.1 or -0.01), reward (+1) or punishment (-1) signal (scores), depending on whether the motor command moved the racket in a favorable direction, or resulted in scoring (hitting the ball) or losing a point (missing the ball).

Visual stimuli (pixel intensities) from the game environment activated a 2D array of time-varying Poisson inputs (20Hz for the feedforward and 35 Hz for the recurrent model) representing the retina in a topographical manner, where the Poisson firing rate was controlled in an all or none manner. These retinal inputs projected topographically to V1. Before driving retinal inputs with pixel intensities, we converted the red/green/blue values to binary values and then down-sampled to a fixed width (80 pixels) and height (80 pixels) to allow all games to provide the same amount of visual information to the model.

The biological visual-motor cortical circuit contains a large variety of neurons, which encode different visual features like location, time, direction, speed, and velocity. We included only location and direction encoding neurons. A population of location encoding neurons received the inputs from the Poisson processes driven by the pixel intensities in a topographic manner, whereas, 8 populations of direction encoding neurons/ direction selective neurons: V1DE, V1DNE, V1DN, V1DNW, V1DW, V1DSW, V1DS, V1DSE (V1 denotes visual area V1, D denotes direction selective neurons, following 1 or 2 letters denote the direction e.g. E for east, W for west, NE for north-east) received inputs from 8 two-dimensional arrays of time-varying Poisson inputs also in a topological manner, where the Poisson firing rates were controlled by the angle of object motion/trajectory. Direction vectors were computed for each object in the visual scene by tracking the position of the object between the last 2 consecutive image frames of the game and only Poisson processes at the location of those positions were driven at particular firing rates.

### Architecture Design Considerations

Our modeling was oriented towards capturing functional aspects of visual and motor areas. We limited visual areas to the minimum needed to represent important game information (location and motion direction of objects, as described above). We used a similar design strategy for the motor area, using a single layer that could make the up/down move decisions. For the association area, we hypothesized that two association layers would allow integration of both object location and movement direction to enable correct motor decisions.

### Initializing weights of synaptic connections

We adjusted initial synaptic weights manually. In this process, the number of synaptic inputs and their average starting weights on specific circuit pathways were adjusted to prevent both hyperexcitability and overly sparse firing. In this way, the later learning would allow increasing or decreasing synaptic weights without totally suppressing neuronal activation, or producing eplieptic-like activation. This allowed all visual, association (premotor) and motor areas to show stable firing rates in physiological ranges.

### Training the models and evaluating learning performance

All models were trained using training episodes, where each episode was simulated for 500 sec while the model was controlling the racket learning to hit the ball. At the end of each training episode, the plastic weights of synaptic connections were saved so that those weights could be used to train the model for the next episodes. The ball and racket positions, as well as the network states, were reset at the beginning of each training episode. To evaluate the learning performance of the model, we ran simulations with the model playing the “Racket-ball” game using the fixed synaptic weights which we selected based on the cumulative hit to miss ratio during training. Each simulation was repeated multiple (6-9) times using fixed synaptic weights and only changing the initial position of the ball and the racket in the court. The performance of the model using fixed weights after training was compared with the model’s performance using the initial random weights to quantify how much the model had learned. All the simulation parameters were reinitialized at the beginning of each subsequent simulation/episode except the weights of synaptic connections.

### Simulations

The model was developed using parallel NEURON (<u>neuron.yale.edu</u>)[58] and NetPyNE (www.netpyne.org) [59], a Python package to facilitate the development of biological neuronal networks in the NEURON simulator. The full source code is available on github (https://github.com/NathanKlineInstitute/SMARTAgent) and ModelDB (https://senselab.med.yale.edu/modeldb/). All simulations were run using MPI on the Linux operating system using Intel Xeon Platinum 8268 2.9 GHz CPUS. Parallelized across 30 cores, 500 sec of simulation time took between 3-6 hours to run, depending on the particular model, and whether we were running with the learning turned on or off for performance evaluation.

## Results

### Constructing a spiking neuronal network model of visual-motor cortex

To test the capabilities of detailed neuronal network circuit models in achieving high performance, we first designed a feedforward model of the visual-motor cortex with visual, motor and association areas each represented as a single layer of spiking neurons (**Figure 1A)**. We connected the neurons across cortical areas only in a feed-forward manner. In the model of visual cortex, we used two functional types of neurons, EV1 neurons encoding location of the objects in the visual field and EVD neurons encoding object motion directions. To make associations between multiple objects in the visual field and their motion trajectories, we included two layers of association neurons, EA neurons and EA2 neurons, where EA neurons were activated by both EV1 and EVD neurons and EA2 neurons were activated in turn by EA neurons.

In this model, we assigned high densities of synaptic connections between visual and association areas (see details in **Materials and Methods** and **Table 2**) so that during learning weakening and strengthening of synaptic weights would shape sparser connectivity patterns. These counterbalancing effects of increasing and decreasing synaptic weights also contributed to network stability. Motor cortex consisted of two neuronal populations, where each population contributed to a specific motor action. We adjusted synaptic weights and connection probabilities to make sure that the visual inputs evoked responses in visual cortex neurons and reliably propagated throughout the neural circuit (see raster plot in **Figure 1B**), finally generating motor commands. The motor commands were generated by comparing the firing rates of the motor cortex EMUP and EMDOWN neuronal populations at each timestep, in a winner-take-all fashion (e.g. when the EMUP population firing rate was higher than that of EMDOWN, a move ‘Up’ motor command was produced, and vice-versa; when firing rates of both populations were the same, the racket was held stationary).

### Tuning learning parameters for reinforcement learning

To learn any visuo-motor behavior, the model must first decode and interpret the visual scene, developing associations between the objects in the scene to understand the visual environment. We could have used unsupervised learning mechanisms for learning spatio-temporal visual associations, but because of the long time scales of unsupervised learning in biology we decided to keep weights of synaptic connections between visual and association areas fixed in the hope that a visual scene including only a bouncing ball and moving racket would not require any plasticity in the early visual areas.

To learn which motor actions must be taken at any instance in a dynamically changing visual environment, the model should first explore the action space by taking random actions under the supervision of a critic, which tells the model the value of an action it recently took in that particular scenario. Such a learning mechanism where the strengthening or weakening of synapses is associated with a critic’s reward or punishment fits within the framework of reinforcement learning (RL). To use RL, we had to deal with two important issues associated with the distal-credit assignment problem, while learning how to play the bouncing ball game 1. The reward or punishment is given after many executed actions, which requires tracking all those actions and all the neurons/synapses contributing to the generation of those actions. 2. We also need to know how recently the neurons/synapses were activated relative to the reward/punishment in order to assign them the correct credit. These issues are tackled in ANNs by recording all of the states, actions, and rewards throughout an episode and then retroactively adjusting the ANN’s action probabilities using discounted episodic returns and backpropagation [34,37,38,60]. Such a replay and update strategy is quite successful in producing a reward-maximizing strategy over a large number of iterations. There is also biological evidence of such learning mechanisms in the hippocampus [61], and could be useful in navigation tasks where an agent could pause to allow replaying the novel sequences. Even that would require adding another SNN modality representing the hippocampus. Furthermore, any phenomenological implementation of such mechanisms with learning on a trial-by-trial basis would be extremely difficult in SNNs.

Therefore in our SNN we used a STDP-RL rule to tackle the credit assignment problem [13,14,40,45,62]. When pre- and postsynaptic neurons both fired within a short time interval, we tagged the synapse between those neurons with an eligibility trace (**Figure 2A**). We can choose time constants for the eligibility trace to stay active depending on how far in time we want to associate the activity of the neuron pair, with the action produced, and the resulting reward/punishment. For distal credit assignment problems in small networks, activation of eligibility traces for long durations may decrease accuracy of the credit assignment due to spatio-temporal cross talk, resulting in the development of nonspecific visual-motor action maps.

Before we tested the standard STDP-RL we proposed another framework based on the intermediate rewards paradigm (IRP; **Figure 2B**) that required prediction of projected ball location for possible hits or misses. Once the model was provided information about the projected ball location, each action delivered a reward when the racket moved towards the target location or a punishment when the racket moved away from the target location. Because each intermediate reward was associated with the past action, we chose a very short time constant (50 ms) for the eligibility trace while using IRP with the feedforward model. When the correct associations between visual space and motor space were established, the model *knew* about the correct action for each visual scene.

Following the standard, non-targeted STDP-RL, all motor neurons become eligible for potentiation or depression based on their spike times relative to the spike times of their presynaptic neurons (**Figure 2C**). Here, we limited action associated rewards and punishments only to the action associated connections and provided opposite and attenuated reward or punishment to non-action associated areas, similar to asymmetric values used in earlier models [63]. For example, if the reward was associated with a “Move-Up” command, the synapses onto EMUP became eligible for potentiation and the synapses onto EMDOWN were made eligible for depression (*targeted RL*; **Figure 2C**). For the plastic synapses not directly making connections onto motor areas, an additional rule (*retrograde targeted RL*; **Figure 2C**) was devised where the reward and punishment were scaled down as a function of the number of synapses away from the motor areas. *Non-targeted RL* and *targeted RL* were used for training feedforward models, whereas *Non-targeted RL* and *retrograde targeted RL* were used for training recurrent models that will be discussed in the later part of the manuscript.

### Training the feedforward SNN to play the racket-ball game

Using a custom built racket-ball game environment (see **Materials and Methods** for details), we first trained our feedforward models with *non-targeted* and *targeted RL* paradigms using both *intermediate* and *sparse* rewards to hit the ball bouncing around the court using the model-controlled racket. The racket movements were generated by comparing the firing rates of neurons in motor areas each representing a different motor action (“Move-up” and “Move-down”). The motor neurons primarily received inputs from the neurons encoding visual features such as the location and motion direction of the objects in the visual scene, and associations between those features in the continuously adapting visual environment. When the model-generated action resulted in a hit or movement towards the ball-projectile, a reward signal was delivered allowing the model to learn associations between the features of the visual space and appropriate actions through STDP-RL. Similarly, when the model-generated-action resulted in a miss or movement away from the ball-projectile, a punishment signal was delivered to weaken the connection weights mediating the associated visual-motor behavior. We chose a smaller multiplicative factor for the punishment and larger multiplicative factor for the reward which caused the weights of the plastic synaptic connections and the firing rates of neuronal populations to increase with training (**Figure 1C**) but in general remained stable and we did not observe depolarization-block anywhere in the circuit during and after training. Only results from the feedforward model utilizing *targeted RL* paradigm are shown in the following sections because our feedforward model utilizing *non-targeted RL* did not learn to play the game.

### Evolution of neuronal circuit properties during training

To investigate how the training affected the dynamics of the modeled neuronal circuit, we first looked at the firing rates of the neuronal populations whose synaptic inputs were allowed to evolve during the training. For the feedforward model, we only looked at the firing rates of EMUP and EMDOWN populations. Since the inputs were discretized over time, analyzing firing rates could be affected by the choice of temporal window size. To avoid that problem, we computed the population mean firing rate sampled over spatially segregated ball trajectories and plotted it against the individual trajectories as experienced by the model during training (**Figure 1C**).

This increase in firing rates resulted due to increase in the synaptic weights of the connections onto EMUP and EMDOWN neurons as the average weight change of these populations is shown in **Figure 1D**. In the feedforward model, both EMUP and EMDOWN neurons showed large variance in the average firing rates throughout the training. During the early training period, the average firing rates varied in the range of ∼0.1-12 Hz, with the mean averaged over the first 10 ball trajectories to be 8Hz. With training, the spread of average firing rates increased to ∼2-30Hz, with the mean averaged over the last 10 ball trajectories to be 20Hz. The net increase in average weights of EMUP and EMDOWN neurons was about 30%.

### Evaluating the performance during training

We trained the feedforward model to play the racket-ball game in episodes, where each training episode was simulated for 500 sec (**Figure 3**). Using feedforward models with different parameters, each time we simulated at least 20 training episodes. Every subsequent training episode resumed learning using the weights of synaptic connections from the end of the previous training episode. This way, the model remembered what it learned during all previous training episodes. For each training episode, we evaluated the performance of the model by taking the ratio of the total number of hits to the total number of misses. The model learned how to play the game, demonstrated through its performance improving strikingly over repeated training episodes (**Figure 3A**): the number of hits increased while the number of misses decreased (**Figure 3B**). However, when we looked at the temporal evolution of performance for training episodes 18 and 19 in **Figure 3C,D and Supplementary Movie 1,2**, we noticed an evolving cumulative hit to miss ratio. The model performed extremely well at the beginning of each of these training episodes (1.78 and 3 for training episodes 18 and 19 respectively), and then the performance decayed before stabilizing at a high level (0.94 and 0.76 for training episodes 18 and 19 respectively).

**Figure 3.**
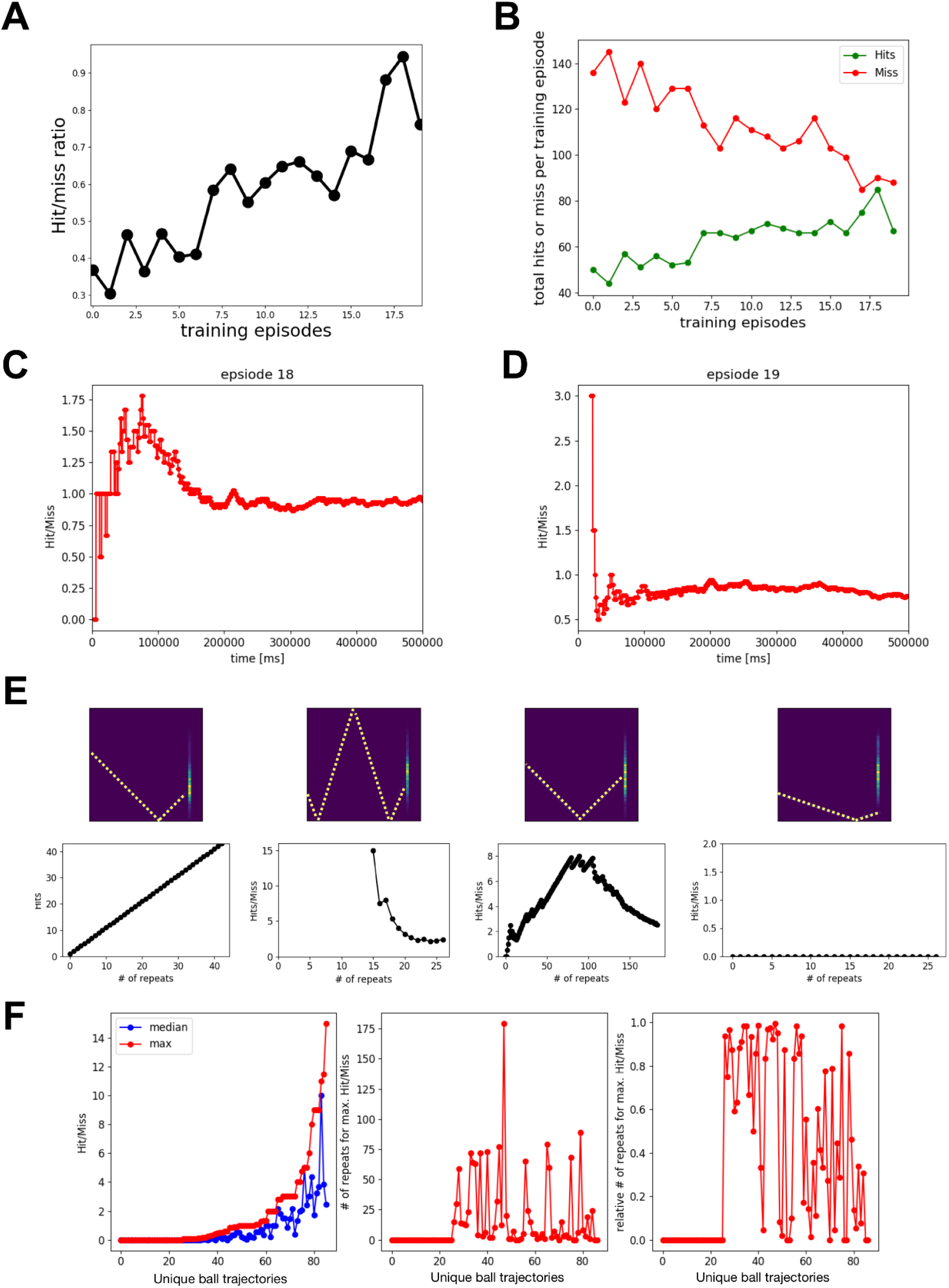
The performance of the feedforward spiking neuronal network model using spike-timing dependent RL improved over repeated training episodes. A) The cumulative Hit/Miss ratio at the end of each 500 sec training episode is plotted as a function of training episodes. B) The total number of Hits and Miss at the end of each training episode is plotted as a function of training episodes. C, D) The temporal evolution of performance for the training episodes 18 and 19. E, F) Summary of learning performance for different ball trajectories. E) Four example ball trajectories are shown together with the performance over repeats. The upper panels show the average of all Input Images corresponding to a unique ball trajectory and the performance is shown in lower panels. These example ball trajectories show visual input specific model learning. For some ball trajectories (e.g. first example), the model-controlled-racket always hits the ball, whereas for some other ball trajectories (e.g. fourth example), it never hits the ball. In the second example, the model-controlled-racket missed the ball only after 15 repetitions. In the third example, the performance first improved, followed by a big drop. F) The left panel: median and maximum performance for unique ball trajectories. The middle panel: number of repeats at which the model had peak performance. The right panel: relative number of repeats at which the model had peak performance. This indicates that for some ball trajectories (# 30-32), the model performed at peak without any training and the training only reduced the performance of the model. For some ball trajectories (# 0-5), the model could not learn to hit the ball. This also shows that for some ball trajectories (see the trajectories with relative # of repeats for max. Hit/Miss values between 0.2 and 0.8), the model first learns to hit the ball and then forgets, whereas for a few ball trajectories (see the trajectories with relative # of repeats for max. Hit/Miss values 0.8 or above), the model did not forget how to hit the ball until the end of all training sessions.

This raises the question of why the performance decreased during training for certain episodes. As the model started playing the game, sometimes the model-controlled racket hit the ball and other times the model-controlled racket missed the ball. The model-controlled racket could hit the ball for three reasons: 1) It learned about the ball trajectory and associated racket behavior, 2) the behavior was intrinsically encoded in the circuit, or 3) completely randomly.

Another important factor to understand the temporal evolution of performance is the varying ball trajectory during the game. The ball could traverse a different path every time it was hit or missed by the racket. So in addition to the three factors explained above, an unseen or unlearned ball trajectory could also explain a decrease in the performance during training.

However, given enough time the model should eventually learn about the new ball trajectories. This line of reasoning motivated us to further dissect the performance of the game based on the ball trajectories.

Using data from all 20 training episodes, we first identified all unique ball trajectories (86) which were repeated at least 5 times. Then for each of those 86 unique ball trajectories, we extracted the hit to miss ratio for all repeats in order of their occurrence. We noticed diverse model behaviors for different ball trajectories that could be explained by learning, intrinsic circuit dynamics, and randomness. We considered each ball trajectory beginning from the time point when the ball started moving towards the racket until it hit or missed the racket. Some representative examples of ball trajectories and the associated performances are shown in **Figure 3E**. The first example ball trajectory (extreme left plot in **Figure 3E)** occurred 43 times and surprisingly the racket never missed the ball. The second example ball trajectory in **Figure 3E** occurred 27 times. Similar to the previous example, the racket always hit the ball during the first 15 occurrences, and only after that the racket missed, bringing the hit to miss ratio to ∼2.

Both of these examples lack any direct evidence of learning as the model never missed the ball, at least for quite a few repetitions. Such performance could be attributed to the intrinsic circuit dynamics emerging from synaptic connectivity patterns and initial synaptic weights.

Unlike the first two presented examples, the model clearly learned about the ball trajectory with repetitive occurrences in the third example in **Figure 3E**, where the hit to miss ratio increased from 0 to 8 over the first 80 repeats, and was sustained around 8 for the following 25 repeats, and only then decreased to ∼2.5 over the next 80 repeats. A few factors that could explain this decrease in performance are: 1) Overlapped visual-motor association, 2) Forgetting, or 3) Lack of association between ball trajectory and some racket positions, since in our analysis we only considered unique ball trajectories and did not control for the racket positions. The first and second factors are not mutually exclusive and are extensively being investigated by the modeling community [64].

Surprisingly, the model could never learn to hit the ball for a few ball trajectories despite many repetitions (e.g. see right most panel in **Figure 3E**). Overall, we found that for 26 out of 86 ball trajectories, the model could not learn to hit the ball. Most of these ball trajectories were targeted towards the corners of the court. However, we could not establish any causal link of this spatial effect to our model’s circuit features or dynamics. For most of the remaining 60 ball trajectories, moderate learning was observed (**Figure 3F)**. For 22 of the remaining 60 ball trajectories, the model’s performance primarily remained improving during the first 80% of the repeats (see red dots above 0.8 in the right panel of **Figure 3F**), whereas for the other 19 ball trajectories, the models’ performance primarily remained declining during the last 80% of the repeats (see red dots below 0.2 in the right panel of **Figure 3F**). For the 19 ball trajectories, the model first learned to hit the ball and then unlearned or kept forgetting (see red dots between 0.2 and 0.8 in the right panel of **Figure 3F)**. Although the model showed peak hit to miss ratios of 15 and ∼11.5 for two different ball trajectories, the hit to miss ratio rapidly dropped to 2 during the later repeats. We found 10 ball trajectories for which the hit to miss ratio remained above 2 after learning. Some other noticeable performances included a ball trajectory for which the model encountered the trajectory 42 times and hit 42 times, and for two other trajectories, the model missed only once after 5 repeats and only twice after 10 repeats.

### Comparing performance of the model after learning with before learning

In the previous section, we presented the performance of our feedforward model during training and noticed some drops in the performance for two cases: 1) when we looked at the cumulative performance during training episodes 18 and 19; 2) when we looked at performance for individual ball trajectories. If there was spatiotemporal interference due to dynamically changing ball trajectories during training, then the cumulative hit to miss ratio might not indicate real performance. Another issue with cumulative hit to miss ratio during training is that it is tracking performance of continuously evolving network states. The ideal test would be to take a snapshot of weight matrices representing a network state and test the model’s performance using those fixed weights without additional plasticity. Furthermore, the performance must be tested against the initial network state to judge how much the model has learned. Next, we addressed some of these issues.

We first simulated our model using initial weights with STDP-RL turned off. To introduce diversity in the ball trajectories, we ran six simulations each with a different initial position of ball and racket and analyzed the cumulative hit to miss ratio for each simulation. We expected the performance of these simulations to differ from one another because as we mentioned earlier the performance depends on the ball trajectories. The hit to miss ratio of 6 simulations with initial weights before training varied between 0.3 and 0.42 with an average of 0.35 (**Figure 4A**). When the simulations were repeated with weights of synaptic connections at the end of the training episode 18, the cumulative hit to miss ratio was substantially larger, varying between 0.72 and 0.89 with an average of 0.8 (**Figure 4A**). Similarly, the weights from training episode 19 yielded improved performance between 0.63 and 0.8 with an average of 0.7 (**Figure 4A**). Note that the performances at the end of training episodes 18 and 19 were 0.94 and 0.76 (**Figure 3C, D and Supplementary Movie 1,2**), which were slightly higher than the respective average performances after training. However, such small differences could be easily explained by differences in the ball trajectories experienced by the model as depicted in the temporal evolution of cumulative performance for simulations before training (**Figure 4B and Supplementary Movie 3**) and after training episodes 18 (**Figure 4C and Supplementary Movie 4**) and 19 (**Figure 4D and Supplementary Movie 5**). Overall, the comparison (**Figure 4A**) clearly showed that the model robustly learned the behavior.

**Figure 4.**
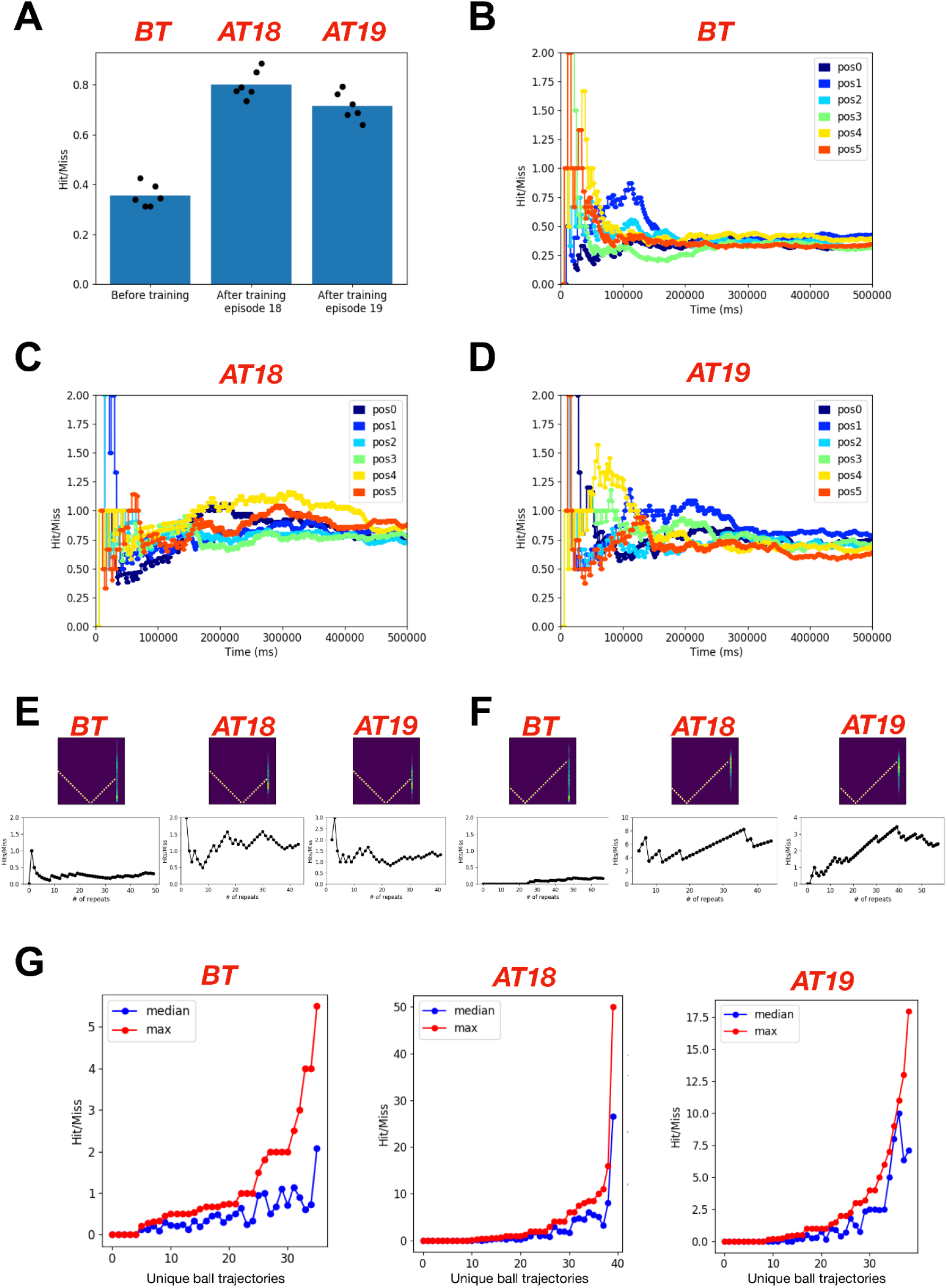
The feedforward spiking neuronal network model sustained its performance after learning. **A)** The bar plot shows the mean (n=6) performance (Hit/Miss) of the model before training (using initial weights), after training episode 18 and after training episode 19. For each condition, 6 different initial positions of the racket and the ball were used to evaluate and compare the performance of the model after learning. Each simulation was run for the duration of 500 sec. **B)** The temporal evolution of the cumulative performance (Hit/Miss) for the model before learning (using initial weights for synaptic connections). The traces in different colors show performance for different initial positions of the ball and the racket. **C)** Same as in **B** using fixed weights for synaptic connections after training episode 18. **D)** Same as in **C** using fixed weights for synaptic connections after training episode 19. **E, F)** Two example ball trajectories where the model showed robust and sustained learning after training episodes 18 (middle) and 19 (right) as compared to before learning (left) **G)** The peak (best cumulative Hit/Miss during repeats) and the median (median of cumulative Hit/Miss during repeats) performance for all different ball trajectories is summarized for the model before training (left) and after training episodes 18 (middle) and 19 (right).

To further investigate the sustained learning of the model, we next compared hit to miss ratios based on the ball trajectories and found that the model showed better performance for most of the ball trajectories after training (**Figure 4G**). Two such examples are shown in **Figure 4E,F**, where the model’s performance was well below 1 before training and increased to much higher value (at least greater than 1) after training. There were still a few ball trajectories for which the model could never hit the ball before and after training (**Figure 4G**).

### Spatial effects of visual environment on learning behavior

As mentioned above, ball trajectories for which the model was unable to ever hit the ball ended up near the edges of the court. We wanted to understand if there were any common spatial features of the ball trajectories, which had an effect on the learning capabilities of the model. Therefore, we classified the coordinates of the ball at the time of reward or punishment into two groups, 1) when the racket successfully hit the ball and 2) when the racket failed to hit (or missed) the ball, and plotted the counts of hit and miss for different y-positions of the ball (at the time of reward; left panels of **Figures 5A,C,E**). The skewed distribution of blue and red bars in **Figure 5A,C,E** shows that the ball moved more frequently towards the bottom edge of the court. Accordingly, the propensity of hits was higher towards the bottom edge of the court before training (**Figure 5A**), which became more uniform with higher tendency to hit towards the center of the court during training (**Figure 5C**). The non-uniformity of hits and misses appeared again in the histogram after training (**Figure 5E**), which suggests that the non-uniformity might be related to the limited sampling of ball trajectories. However, higher red bars compared to blue bars during and after training (**Figure 5C,E**) suggest that the model did not effectively learn the behavior associated with the ball trajectories towards the edges of the court. Longer training that includes sampling of these missing ball trajectories could potentially alleviate these issues.

**Figure 5:**
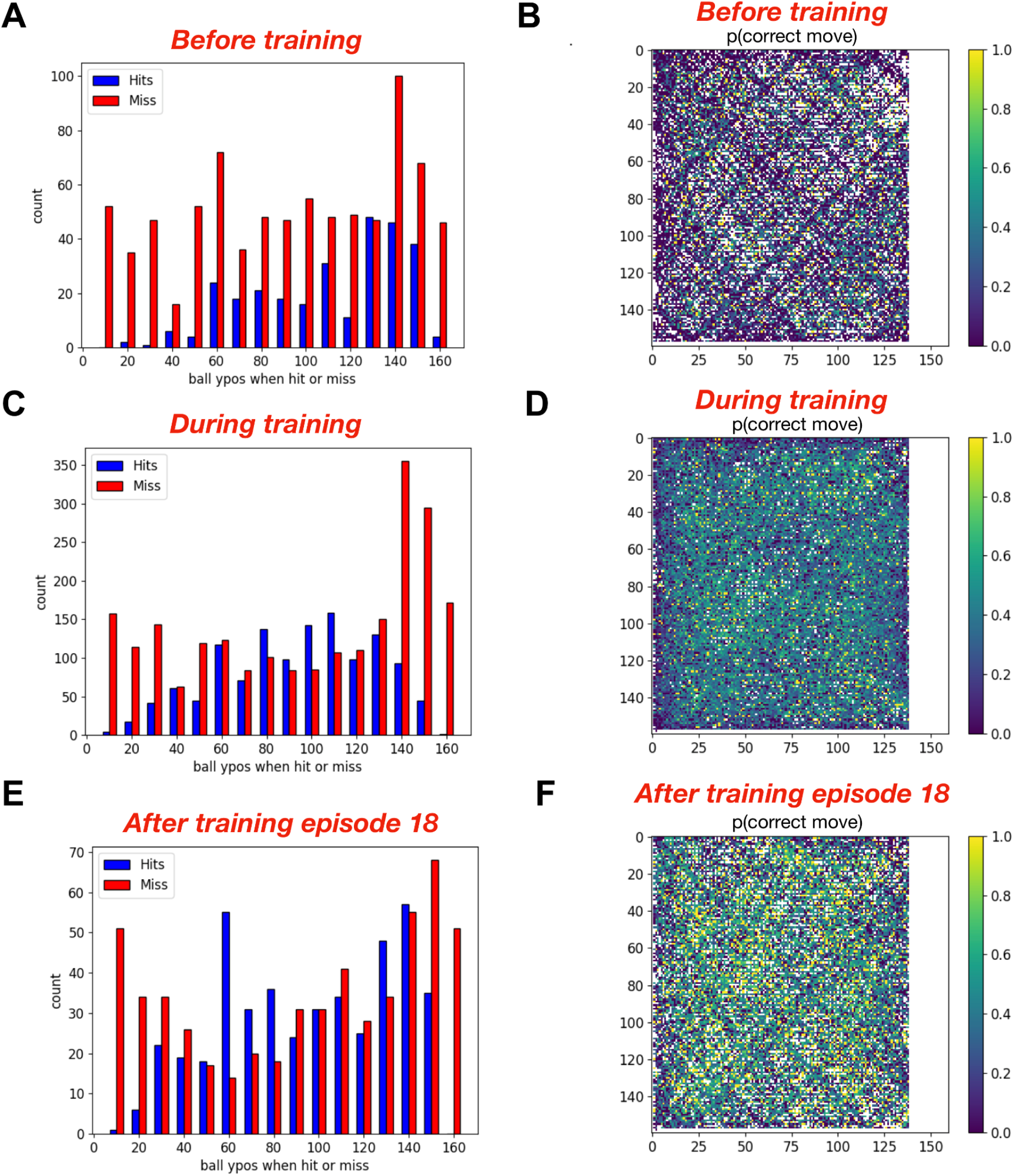
The feedforward spiking neuronal network model learned to perform better for the ball trajectories towards the center compared to the ball trajectories towards the corners of the court. The bar plots in A,C and E show the number of ‘Hits’ and the number of ‘Misses’ against the ball’s vertical position (ypos) when crossing the racket for the model before, during and after training respectively. The heatmaps in B, D and F show the probability of a correct move for each ball location in the court for the model before, during and after training respectively. The color at each pixel in the heatmaps shows the probability of correct action when the ball was at that location based on the projected Hit coordinates (when the action is the same as the proposed action). The white pixels represent the locations never parsed by the ball. Similarly the white space on the right side of each heatmap indicates the region, where no proposed action was available for the model racket (p(correct move) = NaN) as the ball had already passed the racket on the right side of the court.

Although the main training goal was to teach the model to hit the ball, we used intermediate supervisory rewards at all time steps during each ball trajectory moving towards the racket. We used additional cues in our model like projected location of the ball when it would potentially cross the racket and used those cues to teach the model which action was favorable or unfavorable based on whether that action helped the racket reach towards the projected target location or not. To analyze how well the model learned about those cues so that when the ball arrived at each location multiple times, the model produced favorable actions, we plotted the probability of an action generating the correct/favorable move at all traversed pixels of the court as shown in **Figures 5B, D** and **F**. Comparing these probability plots before, during and after training clearly shows that the model was able to identify correct actions at more ball locations after training as compared to before training. Sparser yellow pixels (indicating high probability) in the heatmap during training (**Figure 5D**) might be because of taking into account all data during training, in which case higher probabilities at later training episodes might get masked due to the earlier low probabilities of correct actions.

### Action generation and motor neurons activity

To investigate how persistently and selectively motor neurons get activated during action generation based on the ball trajectories and whether their participation changes after training, we marked all the neurons which were among the top 70% active neurons during each encounter of the ball trajectory. For repetitions of the ball trajectories, we computed the probability of each neuron being among top 70% active neurons and plotted it as a heat map.

The heatmap in **Figure 6A upper panel** shows the probability of each EMUP neuron being among the top 70% most active neurons during repeated ball trajectories before training and the heatmap in **Figure 6A lower panel** shows the same after training (episode 18). Note that the neuron indices in **Figure 6A** are the same but the indices of the unique ball trajectories may differ. Surprisingly some neurons were persistently among the top 70% EMUP population regardless of the ball trajectory (see continuous yellow vertical stripes) and retained such characteristics even after learning (**Figure 6A lower panel**). Some weakly persistent neurons became more persistent after training (see diffusing yellow vertical stripes in **Figure 6A upper panel** becoming solid yellow vertical stripes in **Figure 6A lower panel**), whereas the other weakly persistent neurons consistently knocked out of the top 70% category (see some diffusing yellow vertical stripes in **Figure 6A upper panel** becoming solid blue vertical stripes in **Figure 6A lower panel**). To sum up, the persistently active neurons became more active after training, whereas less persistent neurons, which might be representing the association between visual inputs and respective ‘rewarding’ actions showed two distinct types of behaviors. Some of those weakly persistent neurons became more responsive whereas others became less responsive to the inputs.

**Figure 6.**
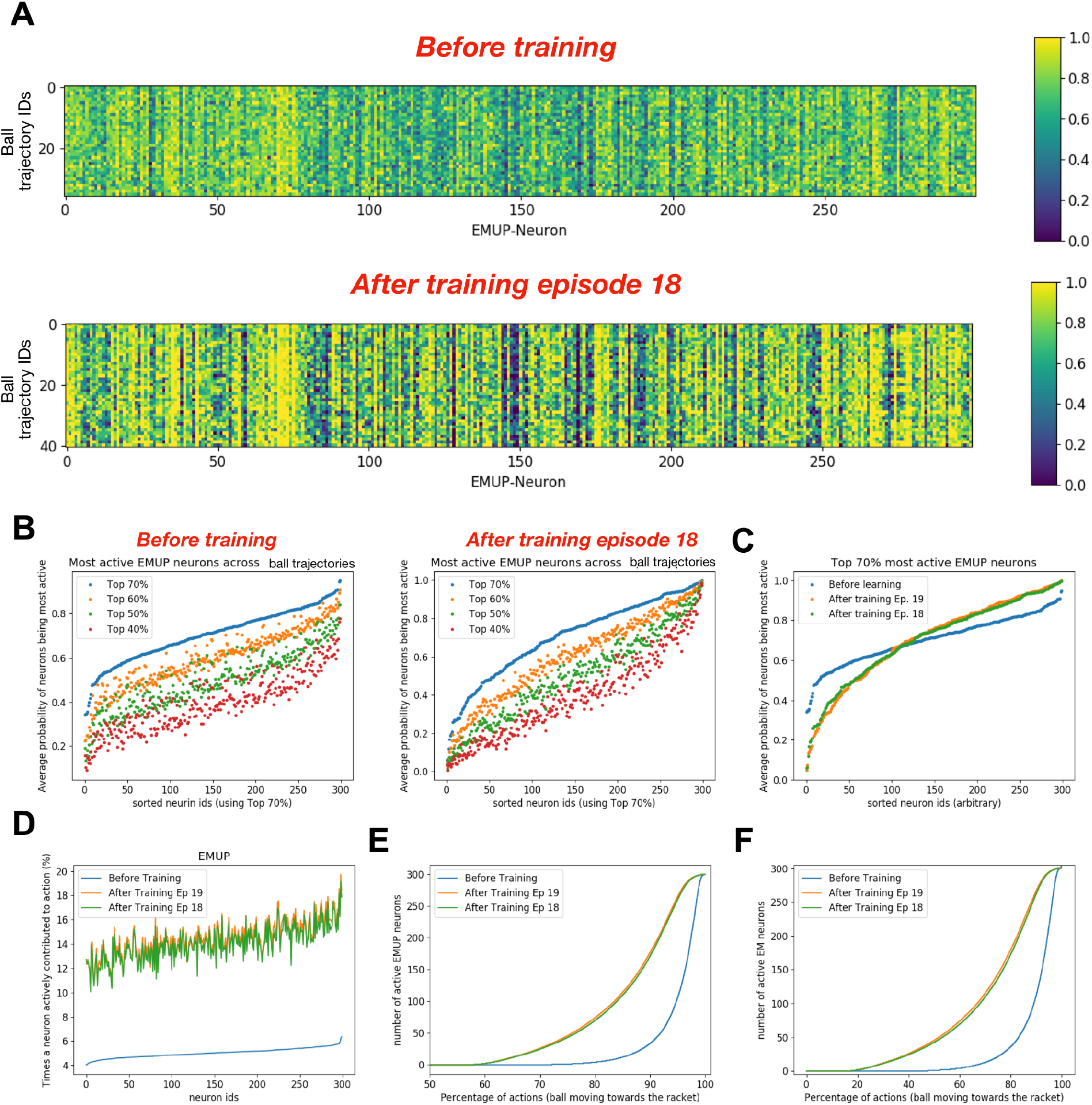
After training, the dynamics of motor neurons taking part in action generation change. **A)** The heat maps show how often each EMUP neuron was among 70% most active neurons during repeated occurance of the same ball trajectory before training (upper heatmap) and after training episode 18 (lower heatmap). Note that the neuron ids are the same in both heatmaps but input seq ids may vary. **B)** The plot shows how often each EMUP neuron was among 70%, 60%, 50% and 40% most active neurons during repeated occurance of the same ball trajectory before training (left: *BT*) and after training episode 18 (right: *AT18*). Note that the neuron ids are sorted using top 70% neuron indices. **C)** Comparing the average probability of each motor (EMUP) neuron being among 70% most active neurons before and after training episodes 18 and 19. **D)** The plot compares the percentage of times a motor (EMUP) neuron actively contributed to action generation. After training, the contribution of each motor (EMUP) neuron in action generation increased proportionally (with some variability) to the contribution before training. **E)** The plot compares how many times at least 1 EMUP neuron was involved in action generation before and after training. Before training, at least 1 EMUP neuron was active for 28% of actions generated. After training, at least 1 EMUP neuron was active for 42% of actions generated. **F)** The plot compares how many times at least 1 motor neuron (either EMUP or EMDOWN) was involved in action generation before and after training. Before training, at least 1 motor neuron was active for 53% of actions generated. After training, at least 1 motor neuron was active for 83% of actions generated.

In the above analysis, a threshold of 70% was chosen arbitrarily, therefore to test whether these observations are independent of the threshold value, we extended our analysis to 40%, 50% and 60% of the most active neurons and plotted the average (across ball trajectories) probability of each neuron being among the top 40, 50 and 60% most active EMUP neurons as shown in **Figure 6B**. High values of average probability could mean either the neuron was persistently among the top X% of the population (where X is 40, 50, 60 and 70) across all ball trajectories, or the neuron was persistently active for the repeated ball trajectories. Similarly, low values of average probability could mean either the neuron was infrequently among the top X% of the population (where X is 40, 50, 60 and 70) across all ball trajectories, or the neuron was sparsely active for the repeated ball trajectories. Note that the neuron identifiers were sorted using the top 70% data. Changing the threshold for data analysis for the simulations before training primarily resulted in linear shifts (**Figure 6B left panel**), suggesting that the relative contribution of each neuron in action generation was uniform e.g. the neuron with largest average probability of being among 70% most active neurons remained the neuron with largest average probability of being among 40% most active neurons. After training, the shifts in the average probabilities of EMUP neurons being among different ranges of activation were nonlinear (**Figure 6B right panel**) indicating non-uniform participation of different neurons.

Furthermore, we found that the neuronal population developed a larger dynamic range after learning (0.32-1; **Figure 6C**) as compared to before learning (0.02-1; **Figure 6C**), indicating better discriminating power of the motor population after training.

Next, we analyzed for what percentage of actions regardless of the ball trajectory, each motor neuron was active. The whole population participated sparsely in action generation before training as the least participating neuron was active only during 4% of actions, whereas the most participating neuron was active only during 6.2% of actions (**Figure 6D**). This increased nonlinearly to 10% participation by the least active neuron and to 20% participation by the most active neuron. The increment in participation of individual neurons in action generation after training was mainly independent of their contribution before training. Even with increased participation of the most active neuron to 20%, many neurons would have to collectively participate at each time point in action generation.

To investigate how many neurons were active during action generation, we analyzed the cumulative probability distributions of active EMUP neurons during action generation (**Figure 6E**). The cumulative probability distribution of active EMUP neurons before training shows that during 72% of actions, no EMUP neuron was active. For the remaining 28% of actions, one or more EMUP neurons were active, with a steep increase in population size of active EMUP neurons during action generation (see blue curve in **Figure 6E**). After training, the percentage of action generation without a single EMUP neuron being active reduced to 58%. For the remaining 42% actions (after training episodes 18 and 19), one or more EMUP neurons were active, with a slower increase in population size of active EMUP neurons during action generation. Does this mean that 72% actions before training and 58% actions after training were generated without motor neuron activity? This is unlikely because these numbers only show EMUP neuronal population’s participation in action generation. When EMUP neurons were silent, EMDown neurons might be actively participating in action generation. To check that, we looked at the cumulative probability distribution of active neurons in both populations of motor areas during action generation (**Figure 6F**). The comparison in **Figure 6F** shows that before training 47% of actions (i.e. No-Move) were generated without any motor neurons being active, whereas after training only 17% of actions (i.e. No-Move) were generated without any motor neurons being active. In this section, we only presented the analysis for the EMUP population, because all the observations described for the EMUP population were consistently present in the EMDown population too.

### Extending the model by incorporating feedback and recurrent connections

The goal of our study was to construct a biologically detailed model of the visual-motor cortex and to train it to learn complex sensory-motor behaviors. As a first step, we successfully constructed a simple version of the model which included only feed forward synaptic connections while ignoring feedback and recurrent connections, which are characteristic of cortical circuitry and are thought to be involved in enhancing learning capacity and computational capabilities of the cortex [20,21,65]. We then trained the simple model to perform while playing a racket-ball game. As a next step in this study, we extended our model by including feedback and recurrent connections as shown in **Figure 7A** (see **Materials and Methods** for details). Expecting that the feedback and recurrent connections would intrinsically inform the circuit about the events back in time and would thus be sufficient to encode motion direction, we therefore excluded direction selective (EVD) neurons from the recurrent model. At the same time, including recurrent connections in the model with plasticity increased the risk of hyperexcitability and depolarization blockade [66, 67]. Therefore, to counteract hyperexcitability and depolarization-blockade, we added inhibitory neurons to the circuit (see details in **Materials and Methods**) in each modeled area. We also added noise inputs to association (EA, IA, IAL, EA2, IA2 and IA2L) and motor neurons (EMUP, EMDOWN, IM and IML) both to maintain minimum firing rates, and also to increase exploration of motor actions and sensory-motor associations.

**Figure 7.**
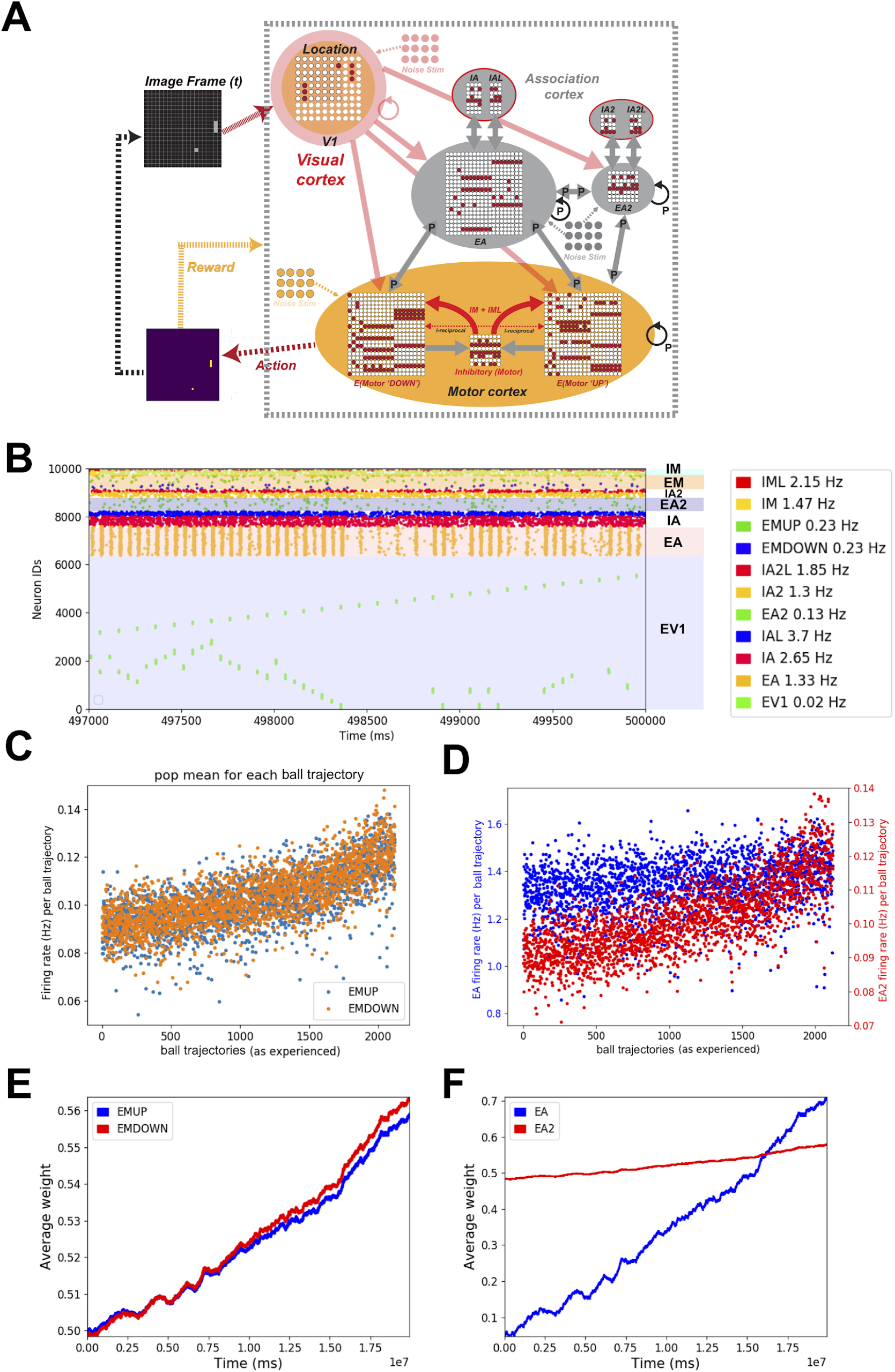
The synaptic weights of the recurrent spiking neuronal network model were adjusted to ensure reliable transmission of the input information across all network areas. A) The schematic shows the racket-ball game interfaced with the recurrent model of visual and motor areas. B) Raster plot showing the spiking activity of different populations of neurons during a training episode. C) Firing rates of motor neuron populations ‘EMUP’ and ‘EMDOWN’ in the recurrent model. D) same as in C for ‘EA’ and ‘EA2’. The firing rates in C and D were binned for ball trajectories (each ball trajectory from the extreme left to the right side of the court where the ball hits or misses the racket). E) Average weight change of synaptic input onto ‘EMUP’ and ‘EMDOWN’ sampled over 40 training episodes. F) same as in E for ‘EA’ and ‘EA2’ sampled over 40 training episodes.

Before training the recurrent model to play racket-ball, we tuned the synaptic weights to allow faithful transmission of spiking activity across the network. Although the addition of recurrent and feedback connectivity increases learning flexibility, it was more difficult to find the parameters that allow learning, while ensuring balanced activity during and after training.

Because of the difficulty to find the appropriateness of the chosen parameters apriori, we trained several versions of the recurrent model with different parameters and evaluated their performance. Although the performance of those models varied, we found that the network dynamics remained relatively stable (see **Figure 7 B-D** and **Supplemental Figure 1A-B**). The firing rates and the weight changes for one example model with good performance are shown in **Figure 7B-D**. An average increase of 12% in the weights of synaptic connections onto EM neurons after 40 training episodes (**Figure 7E**) caused a 30-50% increase in population firing rates. Despite a relatively large increase in firing rates of the EM neurons, the absolute change was minimal i.e. ∼0.03-0.05 Hz (**Figure 7C**). We observed similar characteristics of average synaptic weight (**Figure 7F**) and population firing rate changes of EA2 neurons (**Figure 7D**).

Surprisingly, the population firing rate of EA neurons remained constant during training (**Figure 7D**), despite a 650% increase in the average synaptic weights onto EA neurons (**Figure 7F**).

### Training the recurrent model to learn visuo-motor behavior using sparse rewards

In this study, we showed that using intermediate rewards with a reinforcement learning framework, our SNN models could be trained to perform dynamically adapting visuo-motor behaviors effectively. However, traditionally, sparse rewards are used with a reinforcement learning framework [2, 5]. We next tested the performance of our more biologically detailed recurrent SNN model using reinforcement learning with sparse rewards. To allow association of neuronal activity driving the motor actions with the distal reward, we increased the time constant of eligibility traces to 10 sec. Just like the recurrent model with intermediate rewards (**Supplementary Figure 2A**), the performance increased but kept oscillating between higher and lower values indicating better and worse performance across training sessions (**Figure 8 A, B**). We let the model run for 40 training episodes and found that the model performed reasonably (Hit/Miss = 0.88) well during the training episode 31 (**Figure 8A, B)**. During training episode 31, the model-controlled racket hit the ball 22 times and missed the ball 25 times, which was better than all other training episodes (**Figure 8B**). Overall, the temporal evolution of performance during the training showed similar behavior to the other models i.e. the performance was better during the early training period, dropping to a more sustained value during the late training period (**Figure 8C and Supplementary Movie 6**). As we learned from earlier results that it was difficult to judge learning capabilities of the model during training, we ran control simulations using initial weights and weights at the end of training episode 31. As expected, we observed larger variance in the performance of the model for both cases (**Figure 8D and Supplementary Movie 7**, **8**) i.e. before learning (performance range of 9 simulations: 0.24 - 0.36; **Supplementary Movie 8**) and after learning (performance range of 9 simulations: 0.32 - 0.69; **Supplementary Movie 7**). However, the average performance after learning (0.52) was significantly (p<0.001 using t-test) better than the performance of the model before learning (0.26) as shown in **Figure 8D**.

**Figure 8.**
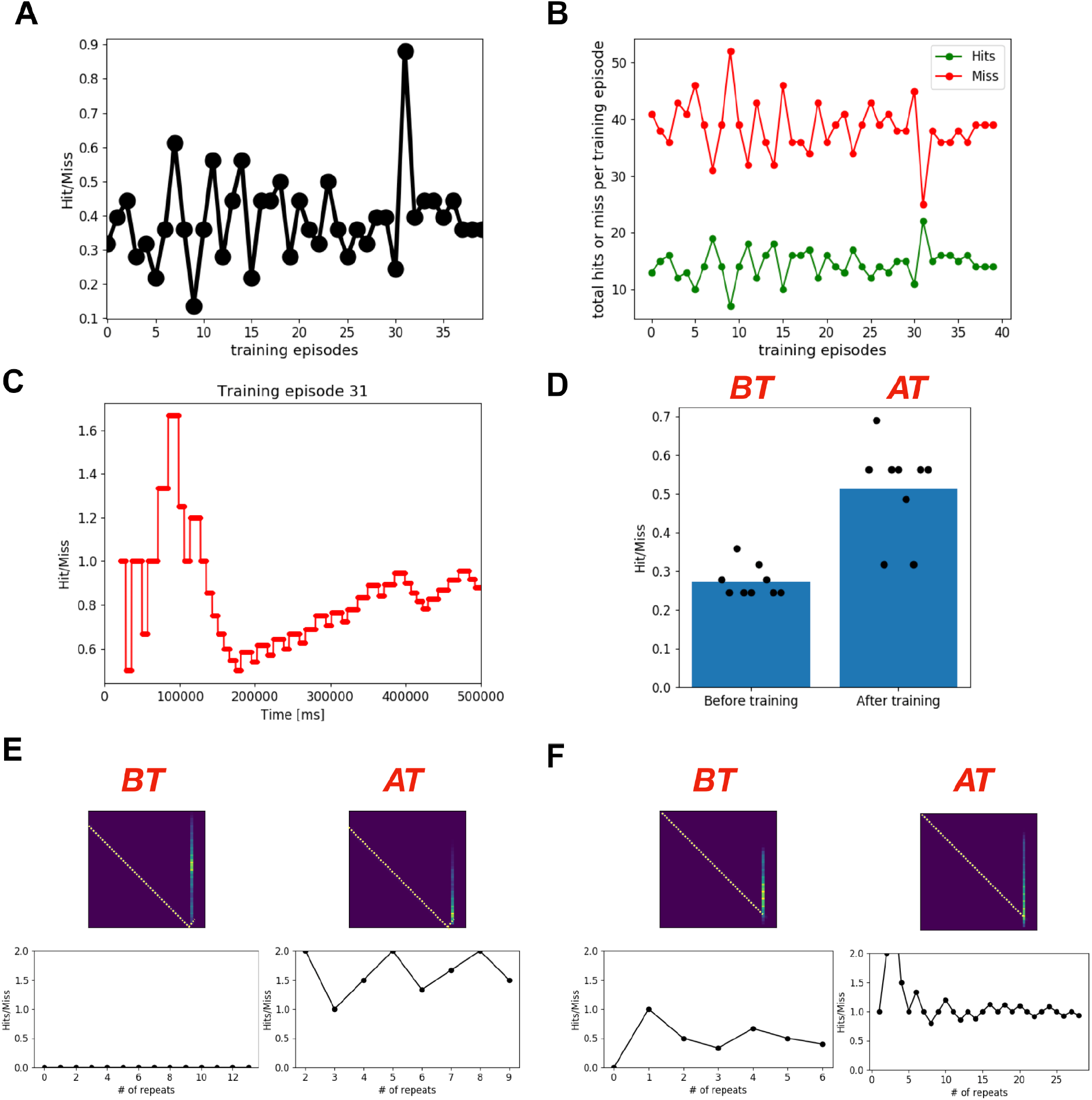
The recurrent model with sparse rewards shows sustained performance after learning. **A)** cumulative performance at the end of 40 training episodes. **B)** Cumulative Hits and Misses at the end of 40 training episodes. **C)** Temporal evolution of performance during training episode 31. **D)** Comparing performance of the model using weights from the end of training episode 31 (right) with the performance of the model before training (using initial weights; left). For both cases, the simulation was repeated 9 times each with different initial positions of the ball and the racket and the performance of each simulation is shown using black dots. The bar plot shows the average of those 9 simulations. **E, F)** Learning by the model is shown using two example ball trajectories. The left panels show the model’s performance for the repeated encounter of the ball trajectory when simulated using the initial synaptic weights (before learning). The right panels show the same as in the left panels but using the synaptic weights at the end of training episode 31 (peak performance *AT* in F is 3).

Next, we compared the performance of the recurrent model before training and after training for individual ball trajectories. Similar to the feedforward model, the recurrent model learned to play the game for many ball trajectories. Two such examples are shown in **Figure 8E**, where the model clearly learned after training. Altogether, these results clearly show that the biologically detailed models with spiking neurons together with reinforcement learning with sparse rewards can be trained to perform complex sensory-motor behaviors.

### Motor neurons sparsely participate in action generation

We had observed a dynamic shift in participation of different neurons in action generation after training the feedforward model (**Fig 6**). We next examined whether those characteristics of neuronal populations persist when we used a more biologically-realistic recurrent model (**Fig 9**). The comparison of heatmaps in **Figure 9A** shows that most of the sparsely active neurons before training did not change their behavior. Instead, they remained sparsely active, likely showing selective responses to the ball trajectories (compare heatmaps for the neuron identifiers greater than 100). Similarly, learning did not have any effect on many of the persistently active neurons (compare yellow colored areas in heatmaps), which were active non-selectively for all ball trajectories. Only a small fraction of the EMUP neurons changed their characteristics after the training as some robustly active neurons before training became more selective to the ball trajectories (compare EMUP-neurons between 20 and 50 in **Figure 9A**). When we changed the threshold for activity participation from 70% to 60%, 50% and 40%, it revealed that the neurons non-uniformly participated in action generation. The least active ∼120 neurons were persistently active during action generation as lowering the threshold did not change the average probability of those neurons being among sparsely active neurons. The other ∼180 neurons were relatively more active but as the threshold decreased their participation probability decreased showing those neurons being selective to the ball trajectories. Surprisingly, these characteristics did not change much after learning (compare panels in **Figure 9B** and **C**) and no increase in discriminability was observed for motor neuronal populations (compare **Figure 9C** with **Figure 6C**).

**Figure 9.**
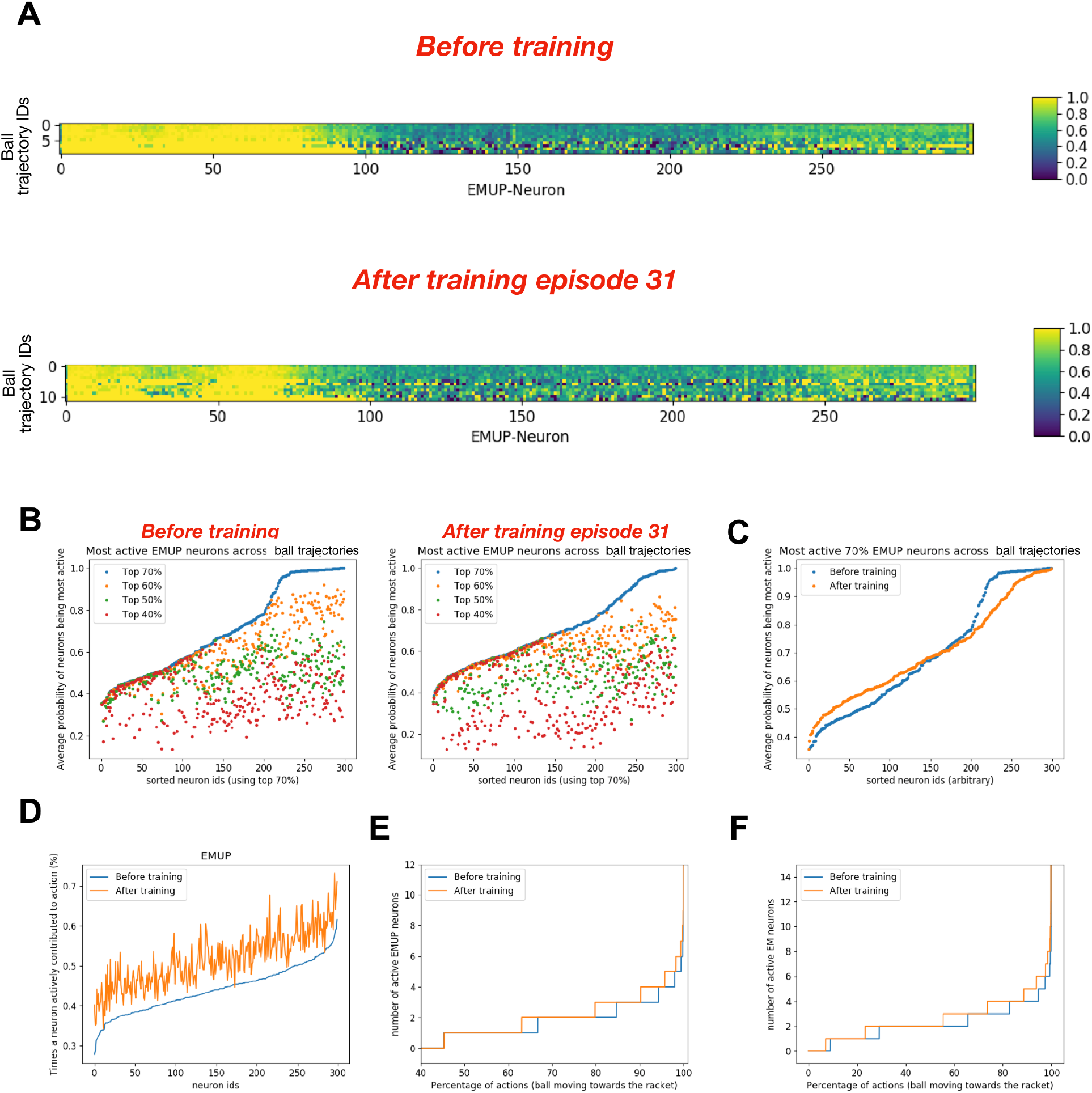
After training the recurrent model with sparse rewards, the dynamics of motor neurons taking part in action generation change. A) The heat maps show how often each EMUP neuron was among 70% most active neurons during repeated occurance of the same ball trajectory before training (upper heatmap) and after training episode 31 (lower heatmap). Note that the neuron ids are the same in both heatmaps but input seq ids may vary. B) The plot shows how often each EMUP neuron was among 70%, 60%, 50% and 40% most active neurons during repeated occurance of the same ball trajectory before training (left) and after training episode 31 (right). Note that the neuron ids are sorted using top 70% neuron indices. C) Comparing the average probability of each motor (EMUP) neuron being among 70% most active neurons before and after training episode31. D) The plot compares the percentage of times a motor (EMUP) neuron actively contributed to action generation. After training, the contribution of each motor (EMUP) neuron in action generation increased proportionally (with some variability) to the contribution before training. E) The plot compares how many times at least 1 EMUP neuron was involved in action generation before and after training. Before and after training, at least 1 EMUP neuron was active for 55% of actions generated. F) The plot compares how many times at least 1 motor neuron (either EMUP or EMDOWN) was involved in action generation before and after training. Before training, at least 1 motor neuron was active for 92% of actions generated. After training, at least 1 motor neuron was active for 93% of actions generated.

Analyzing how actively each motor neuron participated in action generation we found that all neurons were sparsely active during action generation i.e. during less than 1% of generated actions. The least active neuron participated only during 0.28% of actions, whereas the most participating neuron was active only during 0.62% of actions (**Figure 9D**). This increased nonlinearly to 0.35% participation by least active neuron and to 0.74% participation by the most active neuron. The increment in participation of individual neurons in action generation after training was mainly independent of their contribution before training. Overall the relative increase in each neuron’s participation in action generation was much smaller than in the feedforward model. The cumulative probability distribution of active EMUP neurons before training shows that during 45% of actions, no EMUP neuron was active (**Figure 9E**). For the remaining 55% of actions, one or more EMUP neurons were active, with a much less distinctive increase in population size of active EMUP neurons after training (compare blue and orange curves in **Figure 9E**). The percentage of action generation without a single EMUP neuron being active remained the same after training. Although the neurons were sparsely active, either motor neurons, EMUP or EMDOWN were active during 92% of action generations, which slightly changed after training (**Figure 9F**).

## Discussion

In this work, we developed and trained several spiking neuronal network models of the visual-motor cortex to play a racket-ball game using biologically inspired STDP-RL. To train our models, we first proposed two types of reward systems based on *intermediate* and *sparse* rewards/punishments and then evaluated the learning performance of our feedforward (**Figures 1, 3, 4** and **5**) and recurrent models (**Figures 7,8 and Supplementary Figure 1,2 and 3**) using both reward systems in *targeted* and *non-targeted* fashion (**Figure 2**). The goal was to explore the potential of different circuit architectures, connectivity patterns and RL rules in learning visual-motor behaviors. Both feedforward (**Figure 1**) and recurrent (**Figure 7**) architectures facilitated robust learning of the visual-motor behavior except when the feedforward model was trained using *non-targeted RL* and *intermediate* rewards (results not shown). The recurrent model showed better performance using *sparse* rewards and *non-targeted RL* (**Figure 8**) compared to *intermediate* rewards and *targeted RL* (**Supplemental Figure 2**). When we compared the models’ performance after training with before training, we mostly observed a sustained performance. A larger variability in the performance of the recurrent model ( **Figure 8D and Supplemental Figure 3E**) could be attributed to the unattenuated extrinsic noise in the model which was included to allow exploring broader visual-motor associations. When the learning performance was further dissected, we found that the model learned extremely well for most of the ball trajectories (**Figures 3E-F, 4E-G , 8E-F and Supplementary Figure 2D-E and 3G-H**).

Comparing the spiking activity of motor neurons, we found sparser but more sustained activity in the recurrent model as compared to the feedforward model (**Figure 6D-F and 9D-F**). Additionally, we found that all our models recruited more motor neurons in decision making after training (**Figure 6D-F and 9D-F**).

Instead of developing a visual-motor cortex model with detailed anatomical and physiological characteristics, we started with minimal essential details to capture biological realism. We modeled visual (EV1 and EV1D), sensory integration (EA, EA2, IA, IAL, IA2, IA2L), and motor areas (EMUP, EMDOWN, IML, IML2) as a single layer of excitatory and inhibitory neurons. Instead of including dedicated functional neural circuits for object recognition [68–70] and motion direction processing [71, 72], we used standard image processing routines to identify objects and to compute their motion directions. Bypassing the neural processing of thalamocortical circuits of visual processing, we directly simulated the neurons assigned to specific visual features. We set up our models in such a flexible manner that makes it possible to plug-in neural circuits of detailed visual processing later without affecting the functionality of the developed model. The population size of the visual area (80x80 neurons) encoding location was chosen specifically for the visual environment of the game (160x160 pixels). We downsampled the visual field by a factor of 2 to reduce the network size and speed up the simulations. We chose a factor of 2 for scaling down the image because further downsampling introduced additional variability in the evoked responses of the input sensory neurons as it introduced unrealistic changes in the ball and the racket size due to aliasing. For the direction selective neurons (EV1D) we chose smaller populations (400 neurons each), assuming that the varying input from the EV1D neurons onto the EA neurons could be filtered out by more robust input from the location encoding neurons (EV1). For the middle layers EA and EA2, several simulations were run using different population sizes starting from 400 upwards and the final population sizes were chosen based on an increased game performance of the model. Since the transmission of the neural responses across any layer depends on reliability of inputs, the population size of the layer and synaptic weights of the inputs each of the neurons receive, the increased population size of middle layers of the neural circuit should at least exhibit similar performance if the weights are re-tuned properly.

There are many differences between ANNs and SNNs that contribute to differences in dynamics and learning performance. These differences range from differences in how individual neurons are modeled, to circuit architecture and organization. In addition, even the basic learning algorithms differ markedly. At the neuronal level, ANNs typically model neurons as simple linear/nonlinear activation functions, with synaptic connections modeled as integrated scalar weights. In contrast, more biologically-realistic neural circuit models often use biophysically and morphologically detailed neurons, diverse synaptic mechanisms, and detailed microcircuit connectivity patterns. The level of biological realism is often restricted by how much we know about the biophysics of individual neurons, their diversity, their connectivity patterns, and the dynamics of synaptic currents passing through their connections. Even when accurate biological information is available, modeling choices often depend on the computational costs vs benefits of including certain biological features. For example, despite highly nonlinear integration of synaptic inputs in real neuronal dendrites, relatively basic single-compartment neuron models have been shown to capture a reasonable level of neuronal responsiveness [73], which also reduces computational costs significantly. Along similar lines, we used simple integrate-and-fire neuron models with adjusted time constants to reasonably replicate dynamics of real cortical neurons.

For instance, to model synaptic integration properties of real cortical neurons with large apical dendrites, we used longer synaptic delays and time constants for pre-and post-synaptic locations compared to the synapses at the soma as done previously [53]. Although individual spiking neurons may require more computational time than simplified ANN neurons, sparse activation of spiking neurons at the population level (**Figures 1B, 6, 7B and 9**), as shown in biological neural circuits, also reduces computational costs significantly.

The scale and circuit complexity of neocortex can also be used to highlight key differences between ANNs and SNNs. Due to the extremely large number of neurons in the cortex (15 to 32 billion in humans [74]), determining the precise number and type of connections between each pair of neurons is almost impossible. Therefore, connection probabilities are estimated between different sampled neuronal populations to devise connectivity rules. Cortical neurons are organized in a columnar structure with layerwise specific neuronal distributions and complex connectivity rules [11,20,21,75,76]. Capturing those details in silico for accurate circuit reconstruction poses a challenge in itself, let alone using those for tuning to behavior. Based on our simplistic choices for the neuron models and single layer architecture for each modeled area, including biologically constrained connectivity patterns was not a realistic option. Therefore, we tuned connectivity probabilities and initial average weights (**Tables 2 and 3**) by trial and error to choose models that produced excellent performance. However, it is likely that other models with different connectivity patterns would show similar performance after training. Many existing network models either only include feedforward connections or when including recurrent connections, all neurons are fully connected with one another [77, 78]. Both of these configurations contrast with biological circuits where local populations of neurons are often sparsely interconnected [11]. Additionally, none of our models include feedback connections, which are also found widely in biological circuits. Based on common practices and biological realism, we tested both feedforward (**Figure 1**) and recurrent (**Figure 7**) SNN models in this work and both models exhibited good performance after training (**Figures 4A and 8D**).

Training for multiple tasks, where meta learning capabilities are required may show additional advantages of using recurrent and additional feedback connectivity motifs, compared to solely relying on feedforward connections. However, in the models used in this work we did not rely on feedback connections, so have not yet ascertained their benefits. In future work, we aim to disentangle the role of each circuit motif in learning behavior, by perturbing circuit architecture and measuring performance.

Another important theme in neuronal network modeling is the balance between excitation and inhibition, which stems from the types of neurons and synapses used in ANNs and SNNs. In contrast to SNNs, ANNs usually lack exclusively defined excitatory or inhibitory neurons since excitation/inhibition can be modeled by simply using positive/negative weights. In addition, ANNs usually do not need inhibition to achieve stable performance, and including inhibition could even impair their learning abilities. Recently, inspired by feedforward inhibitory interneurons in the brain, inhibition was included in an ANN model without sacrificing its learning performance [79]. In SNNs where the weights may change due to learning, too much excitation to a neuron can either lead to depolarization blockade or hyperexcitability, both of which can prevent a neural circuit from functioning properly. In biological neural circuits, at least two kinds of mechanisms prevent neurons and neural circuits from arriving at such states: inhibition and homeostatic plasticity [66]. In our work, including inhibitory neurons in modeled cortical areas with plastic synaptic connection was sufficient to keep the network output and performance stable. Including homeostatic plasticity rules in our model might allow continued learning in a biologically plausible manner.

The learning algorithms used in ANNs and SNNs also differ in important ways. Learning In feedforward ANNs takes place through a backpropagation algorithm, in which synaptic weights are adjusted proportional to the error gradient which is fed backwards from output to input layers [60]. In addition to biological implausibility, such methods are also computationally expensive as they require repeated ‘playback and update’ sessions. In contrast, strong evidence for plasticity mechanisms such as spike-timing dependent plasticity exist in biological neural networks [80], where the learning takes place locally and on a trial-by-trial basis. Therefore it was a natural choice for us to use STDP based learning mechanisms in our spiking neural networks.

Besides biological plausibility, one obvious advantage of using hebbian plasticity rules such as STDP-RL is that network models learn in real time and do not require training using information from the entire game/task, as often used in nonbiological ANNs. Hebbian plasticity rules are deemed so effective in learning that even when adapted for use with ANNs, they show rapid and robust learning behavior [81]. A potential limiting factor in backpropagation and other optimization based learning strategies is their lack of support for continued learning as the weights are required to be kept fixed after training to maintain robust performance. In biological neural networks, synaptic strengths continuously evolve [82–87] due to homeostatic maintenance and random turnover [88–90], however it remains unclear how memories of learned experiences are maintained given continuously changing synaptic connections. Compensatory plasticity mechanisms have been proposed to account for maintaining the memories in the face of such constant synaptic change [91, 92]. Compensatory plasticity can be induced by external reinforcement signals [93], interactions between different brain areas and circuits [94], or spontaneous, network-level reactivation events [95]. Inspired by these mechanisms, Najarro and Risi further showed added advantage to the hebbian and other biological plasticity mechanisms. Including additional plasticity mechanisms and randomizing synaptic connections and weights might lead to even better performance as such capabilities of multi-timescale learning in SNNs for learning visual-motor behaviors have been shown in another study [96].

Animals learn sensorimotor behaviors primarily through three distinct mechanisms: reinforcement learning via striatum, error-based learning via cerebellum, and use-dependent learning in the cortex [31,97,98]. Ideally all those structures capturing biological details and function should be included in the model of sensory-motor learning. Although a lot is known about individual circuits, how those circuits operate in parallel juggling the tasks remains unclear.

Therefore, integrating those circuits in our model would be almost impossible and probably overly done for the task in hand. Reinforcement learning only tells whether the action performed was “good” or “bad”, primarily relies on a reward, and it does not tell how to improve the accuracy of the action to reach the goal, which is instead learned in the cerebellum utilizing sensory feedback. Furthermore, to learn the behavior only via reinforcement learning would require exploration of many actions to choose the best action leading to the reward and it may take an extremely long time to experience the best action, if ever. Supervised learning via cerebellum helps fine tune motor actions to rapidly produce optimal action. Since we wanted to test effectiveness of learning mechanisms operating at different time scales in a comparable manner, instead of building circuit models tackling different time scales, we designed our learning rules mimicking those timescales e.g. we used *intermediate reward* based on visual feedback with 50 ms eligibility trace capturing the working of error-based learning in a “black-box” and sparse-reward with 10 s eligibility trace allowing integration of many actions to the behavioral outcome (hit or miss the ball). Some might argue that our intermediate reward paradigm is cerebellum-dependent and biologically non-overlapped with RL based circuits, however, even in that case there are clearly interactions between cerebellum and basal ganglia in the brain [99, 100], and our intermediate reward paradigm could be viewed as a phenomenological implementation of this type of interaction. RL based on intermediate reward has been used previously in an arm model, which was trained to reach a fixed target [14]. Here, we extended the RL paradigm to utilize both the *intermediate* and *distal*/*sparse* rewards/punishments (**Figure 2B**). Although we used both the *intermediate* and the *sparse* rewards in the feedforward model, choosing a single small time constant for eligibility traces prevented developing associations between distally active neuron pairs (in time) to the actual reward. Using both *intermediate* and *sparsely* occurring actual rewards with RL would require separate mechanisms with different eligibility time constants (shorter time constant for *intermediate* rewards and longer time constants for *sparsely* occurring actual rewards) in parallel. Alternatively, the brain could use multiple types of rewards for the reinforcement learning [101–105] (e.g. could use different types of dopamine receptors), but we are not aware of any direct experimental evidence of how associations between different rewards and respective motor actions take place at multiple time scales. Combining different timescales of reward/behavior in parallel, or independently [96] has also been shown to enhance performance. The time-scales underlying other learning mechanisms (such as homeostatic plasticity, sleep consolidation, etc) could provide additional benefits [106]. We therefore expect that including credit assignments at multiple time scales in our model will further improve its performance.

In addition to using *intermediate* rewards, we also developed a new *targeted RL* algorithm (**Figure 2C**). Instead of providing reward to all neuronal populations, we provided a reward or a punishment only to the neuronal population responsible for the associated action. While there is no direct evidence from biology supporting our exact implementation of the targeted RL framework, we assume that such mechanisms exist based on the experimental evidence of topologically distinct cortico-striatal loops across sensorimotor areas (Foster *et al.* 2021). Such specific feedback configurations could allow distinct motor areas projecting onto the striatum to release dopamine only in specific subnetworks. Some anatomical evidence from invertebrates suggests nonuniform delivery of reward prediction error signals across brain areas in compartmentalized manner [107]. We took a step further and proposed our *targeted RL* to evaluate its potential as a ***proximal credit assignment*** mechanism. To speed up learning, we also provided asymmetrical reward/punishment to the nonassociated population i.e. if the reward was delivered to the EMUP population because Move-Up was the expected action and the model generated Move-Up command, then some punishment was delivered to the EMDOWN population. The *targeted RL* was essential for the *intermediate* rewards due to their frequent occurrences, otherwise many nonassociated pre and post motor neuronal pairs would have encoded nonselective associations. Some evidence of such selective reward based learning can be found in invertebrates [108], where selectivity is often implemented by anatomical constraints. Furthemore, we proposed ***retrograde targeted RL*** where a scaled down reward or punishment for the neuronal connections not directly synapsing onto motor neurons (if those neurons are involved in reinforcement learning), where the scaling factor could be based on the number of connections between the postsynaptic neuron and motor neurons. Such strategy was based on observed lower distribution of dopamine receptors in early sensory areas (Froudist-Walsh et al. 2021), however the exact value was only chosen to estimate biological variance in dopaminergic signaling across cortex. We found all these strategies to be equally effective in learning behavior (**Figures 4A and 8D**) with differences in temporal evolution of learned behavior (**Figures 3A and 8A**).

However, it must be noted that the goal of training using *sparse* rewards was to maximize the hits, whereas the goal of training using *intermediate* rewards was to move the racket towards the projected ball trajectory which would eventually lead to hitting the ball. This is evident in the heatmaps showing the increased probability of the racket moving towards the projected ball location for the hit after training (compare **Figure 5F with Figure 5B**). Based on our results, we hypothesize that a brief and localized delivery of reward prediction error signal could encode temporally precise associations between sensory information and motor actions. Such a system could work in parallel to the global reward prediction error generating system which is thought to mediate distal credit assignment of rewards to sensory cues and associated actions.

When we used the *non-targeted RL* with the feedforward model, the model could not learn the behavior despite trying different parameters and training for several episodes (results not shown). One possible reason for the feedforward model’s inability to learn the behavior could be the use of both intermediate and sparsely occurring rewards with a single eligibility trace (fixed time constant). The problem might have occurred due to the temporal incompatibility of both types of rewards with a single eligibility trace i.e. intermediate rewards are appropriate for associations between neurons at each step whereas sparse rewards require a memory trace of all steps (mediated via long eligibility traces) leading to the reward generation. The frequently occurring intermediate rewards might have strengthened the synapses between coactive pre and post neuron pairs driven by intrinsic noise and might have interfered with neuronal activity generated in the following steps before the actual reward got delivered, by changing the state of the network many times before the associations between the actual reward and neuronal network were established. However, when we used *non-targeted RL* and *sparse* rewards with the recurrent model, the model learned to hit the ball over repeated training episodes (**Figure 8**). This time the proper associations between the ball trajectories and actions to improve the chance of a hit were made because of the long time constant for eligibility traces (10 sec) which acted as a memory trace for the neuron pairs active during action generation for all steps during the ball trajectory.

Although our model learned to play the bouncing ball game effectively, the performance during training plateaued after some episodes suggesting that the model learned to its capacity despite continued increase in weights. Increase in synaptic weights can encode learning as long as they can differentially activate the postsynaptic neurons. If an increase in weight does not change the firing rate of the postsynaptic neuron, the learning remains ineffective. This could happen for intermediate timesteps when the increase in weights is so small that it does not translate into increase in firing rate but would count towards learning after multiple increments in weights eventually increasing the firing rate. In another scenario, any further increase in weights could push the neuron to depolarization block. An effective strategy to increase the capacity of the network could be homeostatic synaptic scaling which has been shown to significantly enhance the performance of neuronal networks [109, 110]. Several models of synaptic scaling have been developed [66,67,111–113], each with their own advantages and biological plausibility. Inclusion of synaptic scaling in the future model is also necessary to allow learning more behaviors without pushing the network towards hyperexcitability leading to seizures. Additionally, increasing the size of the middle and output neuronal populations in the model could also be helpful in increasing the memory storage capacity of the network.

When we first looked at the temporal evolution of the models’ performance, we could not understand why the performance is better during some training episodes and worse during other training episodes. One possible explanation for such variable performance could be the noise inputs that we included in the models to allow more exploration of the motor actions. Most probably that is the case with the recurrent model as we see a larger spread of performance measurement during (**Figure 8A**) and after training (**Figure 8D**) and in fact a lot more noise inputs were used in the recurrent model as compared to the feedforward model. The plasticity for noise inputs will be included in the future models to address noise induced variability in the performance, so that as the visual-motor associations develop in the circuit, the noise input becomes weaker [114]. But this is not the only factor driving variability in the performance. If we break down the bouncing ball game into individual ball trajectories, we notice that the ball trajectories vary from one training episode to the other. The variety of ball trajectories arises because every time the ball is hit, it moves along a different path depending on the point of impact between the ball and the racket. Depending on which ball trajectories have already been learned by the model, the performance may vary from one episode to another. Therefore to make a fair comparison of performance between different learning states (fixed weight matrix after learning) and naive states (before learning), we compared the performance of the model based on ball trajectories. This analysis further revealed that our models learned very well for some ball trajectories compared to a few others (**Figure 3E and F, Figure 4E,F and G**). For most of the learned ball trajectories, the model showed sustained performance (**Figure 4F**). However, for the others, we observed that the performance decreased with repeated encounters of the same ball trajectories (2nd and 3rd ball trajectories in **Figure 3E**).

The neural activity patterns representing past experiences are replayed widely across hippocampus and cortex [115, 116] during sleep and passive awake states [117–119]. These replays are thought to be crucial in stabilizing existing memories and allowing continual learning of other tasks. In the absence of such mechanisms, the brain may forget what it learned. Despite impressive performance in state of the art ANNs, sometimes old memories are lost upon training for new tasks. Storing and replaying learned sequences of neural activations can rescue the old memories but can be computationally expensive. To overcome this limitation, van de Ven et al. [120] proposed including a reservoir/generator which can learn and store different activation patterns and can mimic replay of different neural sequences randomly. Our SNNs faced similar challenges of forgetting even within a task. This might be linked to the network capacity or architecture choice, which resulted in encoding overlapping sensorimotor representations. More specifically, the decreases in performance after learning could be due to overlapped representation of multiple ball trajectories in visual-motor space so that changes in the circuit required to learn about one ball trajectory may interfere with the visual-motor representation of the other ball trajectory. These issues have been observed in ANNs and are commonly termed “catastrophic forgetting” [64,120–125] and could be mitigated by expanding the dimensions of the circuit and including recurrent computing reservoirs representing PFC and Hippocampus. The observed “forgetting” behavior in our model could also be explained by our choice of RL model, where we used reward to model dopamine **as has been extensively done in earlier biological models of RL** [13,14,16,33,45,109], whereas in vivo dopamine encodes reward prediction error (RPE) [126]. Using reward instead of RPE prevents the model from not learning once the sensory cues become familiar and the model has learned the correct action or sequence of actions that could lead to over-writing of information.

Another feature we observed was that for some ball trajectories, the model always hit the ball from the very first encounter and continued to do so until the end of the simulation (e.g first ball trajectory shown in **Figure 3E**). Something similar is observed for other ball trajectories, where the model *knew* to hit the ball for many encounters and then further training led it to forget. In both cases, the performance profile suggests that the naive state (initial weights) of the model was sufficient to capture the visual-motor association for these particular ball trajectories. It would be interesting to further dissect out the anatomical features of the network that enable such intrinsic performance.

Animals use cognitive maps to relate tasks and goals to relevant rewards. In the absence of neural mechanisms encoding cognitive maps of tasks, ANNs or SNNs require defining the goals or behaviors of the sensorimotor tasks explicitly. Just like a point awarded to the winning player when the opponent fails to hit the ball in tennis, other types of rewards can be explicitly defined and used for the training. More complex tasks like any strategy game or chess, which may require including higher cognitive function areas in the model for devising strategy or planning, are beyond the scope of this work.

Our motivation for this work was to develop a framework that could be further extended to incorporate more biological details in spiking neuronal network models to bridge the gap between responsiveness and functioning of the circuit and its elements. Does our current approach and model allow bridging that gap? In a recent paper [127], the authors demonstrate the similarity between the responsiveness of V1 neurons and receptive fields generated by convolutional neural networks (CNNs). In our work here, we did not implement biophysically detailed visual system features using a layered approach following the canonical model of the visual system (V1→V2→V4→IT), as it would have added unnecessary complexity to our model for the required task. Instead, we designed our SNN to simply capture the functionality of visual cortical areas by driving location and direction selective neurons using topographical information consisting of object location and motion direction. Our goal was to train the network to use this information to learn a visual-motor task using biology based mechanisms. In future work, we plan to increase the biological realism of our models’ visual and motor areas, and explicitly compare neuronal responsiveness with neuronal electrophysiology data.

In conclusion, we developed a framework to train more biologically detailed spiking neuronal network models to perform sensorimotor-dependent behaviors in dynamic visual environments. Moving forward, including detailed circuit elements of cortico-basal ganglia loops and other interacting sensory-motor learning modalities like cerebellum would allow a more detailed exploration of the biological mechanisms of sensorimotor behavior and learning.

Electrophysiological recordings from sensory-motor circuits are scarce in behaving animals but once available the model could be further refined to match both electrophysiology and behavior. Testing the related hypotheses using this framework would require adopting both behavioral paradigms as well as the modeled circuit elements. In addition to circuit elements supporting reinforcement and cerebellar learning, we aim to expand this framework by adding other sensory modalities such as auditory [128] and somatosensory. We will be sharing the software and the model with the neuroscience community to expand its functionality and use in research.

## Acknowledgements

This work was funded by ARO W911NF-19-1-0402, ARO-URAP supplement, NIDCD R01DC012947, NIH U24EB028998, NIBIB U01EB017695, NSF 1904444-1042C, and Google Cloud Platform Research Credits. The views and conclusions contained in this document are those of the authors and should not be interpreted as representing the official policies, either expressed or implied, of the Army Research Office or the U.S. Government. The U.S. Government is authorized to reproduce and distribute reprints for Government purposes notwithstanding any copyright notation herein.

## Supplementary Material

**Supplementary Figure 1:**
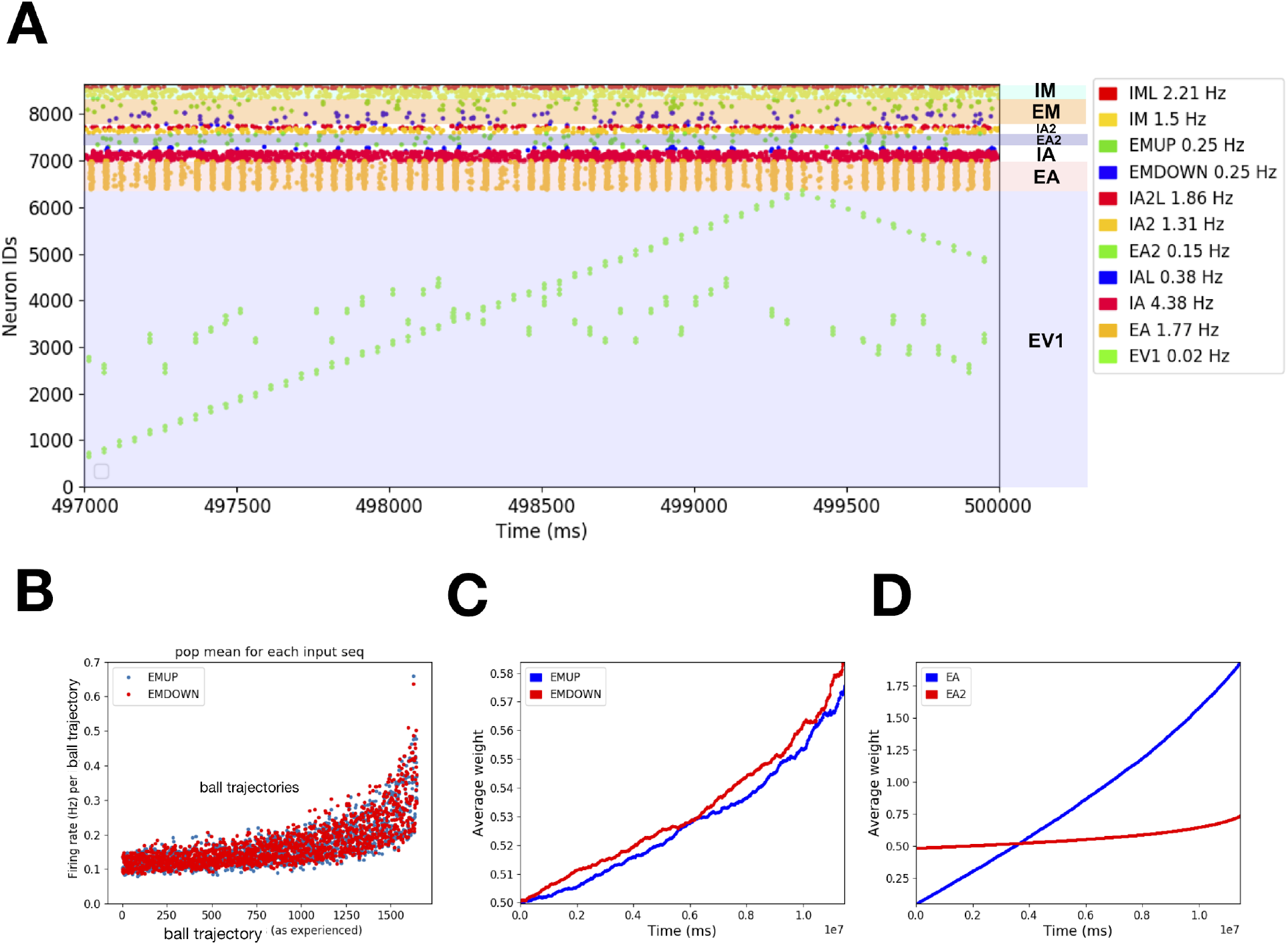
**A)** Raster plot of different populations of neurons during a training episode (vertical axis is neuron identity and horizontal axis is time; each dot represents a single action potential from an individual neuron). **B)** Firing rates of excitatory motor neuron populations EMUP and EMDOWN in the feedforward model increase over the course of training. The firing rates were binned for ball trajectories (beginning when the ball is at the extreme left side of the court and ends when the ball hits or misses the racket on the right side of the court). **C)** Average weight change of synaptic inputs onto EMUP and EMDOWN sampled over 23 training episodes tends to increase with learning, **D)** Same as in **C** for EA and EA2 populations.

### Training the recurrent model to learn visuo-motor behavior using *retrograde targeted RL*

In contrast to the feedforward model, where STDP-RL mechanism was included only at the synapses onto the motor neurons, in the recurrent model, STDP-RL mechanism was also included at the synapses onto the association neurons, EA and EA2. Because these synapses were not directly involved in action generation, we used a special rule for reinforcement learning that we termed as *retrograde targeted RL* in which the synapses away from the motor areas get partial reward or punishment depending on the ‘critic’ value. Next, similar to the feedforward model, we tuned the parameters of the recurrent model to ensure reliable transmission of neural activity across the modeled areas without causing hyperexcitability or depolarization-block (see raster plot in **Supplementary Figure 1A**).

We trained the recurrent network model to play a bouncing ball game for 23 episodes. During training, both EMUP and EMDOWN neurons in the recurrent model were sparsely active (∼0.08-0.15 Hz) at the beginning and later evolved to higher yet still sparse firing rates (∼0.2-0.5 Hz) as shown in **Supplementary Figure 1B**. These firing rates were computed for the duration of full ball trajectories from the left side of the court to the right side and show that during training the model experienced 1600 ball trajectories/ input patterns. The increase in firing rates of motor neurons resulted from increase in the synaptic weights of the connections onto EMUP and EMDOWN neurons as well as increase in the synaptic weights of the recurrent and feedback connections onto EA and EA2 neurons as the average weight change of these populations is shown in **Supplementary Figure 1B**. The net increase in average weights of EMUP and EMDOWN neurons was about 16%, whereas the net increase in average weights of EA and EA2 neurons was 1800% and 40% respectively.

In **Figure 3**, we saw that the performance of the feedforward model clearly improved over repeated training episodes. Although we saw increased performance across the first few training episodes of the recurrent model with IRP (**Supplementary Figure 2A and B**), the performance fluctuated over later training episodes (**Supplementary Figure 2A and B**). Even the best performance (0.6) was not as good as the performance of the feedforward model (0.94), however we clearly observed some learning. We also noticed that the recurrent model learned more rapidly than the feedforward model as the recurrent model’s performance improved to 0.5 only after 4 training episodes (and fewer ball trajectories as each action timestep was 50 ms) as compared to 8 training episodes (and more ball trajectories as each action timestep was 20 ms) for the feedforward model. Most of the performance features during training episodes (**Supplementary Figure 2C-E and Supplementary Movie 9-12**) were similar to what we observed for the feedforward model (**Figure 3C-F**) and are described below.

During 23 episodes of training, overall the recurrent model experienced 46 spatially unique ball trajectories out of which the model could not learn to hit the ball at all for 6 ball trajectories (see last example ball trajectory in **Supplementary Figure 2D**). For 6 of the remaining 40 ball trajectories, the model’s performance (hit to miss ratio) primarily kept improving during the first 80% of the repeats (see red dots above 0.8 in the right panel of **Supplementary Figure 2E** and first and fourth example ball trajectories in **Supplementary Figure 2D**), whereas for the other 18 ball trajectories, the models’ performance primarily kept declining during the last 80% of the repeats (see red dots below 0.2 in the right panel of **Supplementary Figure 2E** and second example ball trajectory in **Supplementary Figure 2D**). We found that for the 16 ball trajectories, the model first learned to hit the ball and then unlearned or kept forgetting as is indicated by red dots between 0.2 and 0.8 in the right panel of **Supplementary Figure 2E** and the third example ball trajectory shown in **Supplementary Figure 2D**. The model displayed an optimal performance for a ball trajectory where the peak hit to miss ratio was 7 and the minimum value for hit to miss ratio (not shown). The best sustained performance was observed for the ball trajectory for which the hit to miss ratio remained around

0.8 for about 80 repeats (see fourth ball trajectory in **Supplementary Figure 2D**). This ball trajectory was repeated most frequently over 120 times during the training episodes. One of the reasons for a large variance in performance during training could be the intrinsic noise in the circuit which was intentionally kept higher to allow the circuit to explore action space to its full capacity. Ideally, the drive by the noise processes should decrease with learning to enable motor neurons to take actions based on the sensory inputs and the sensory-motor associations the model learned during training. However, we have not tested the use of adaptive noise in this work. The other reason for large variance in performance could simply be the fact that the performance presented in **Supplementary Figure 2** is during learning which is an extremely dynamic situation. Since each training episode (or controls) was simulated for 500 sec, using larger action timesteps of 50 ms (compared to 20 ms for feedforward model) reduced the repeats of each ball trajectory which would have caused a large variance in performance.

**Supplemental Figure 2.**
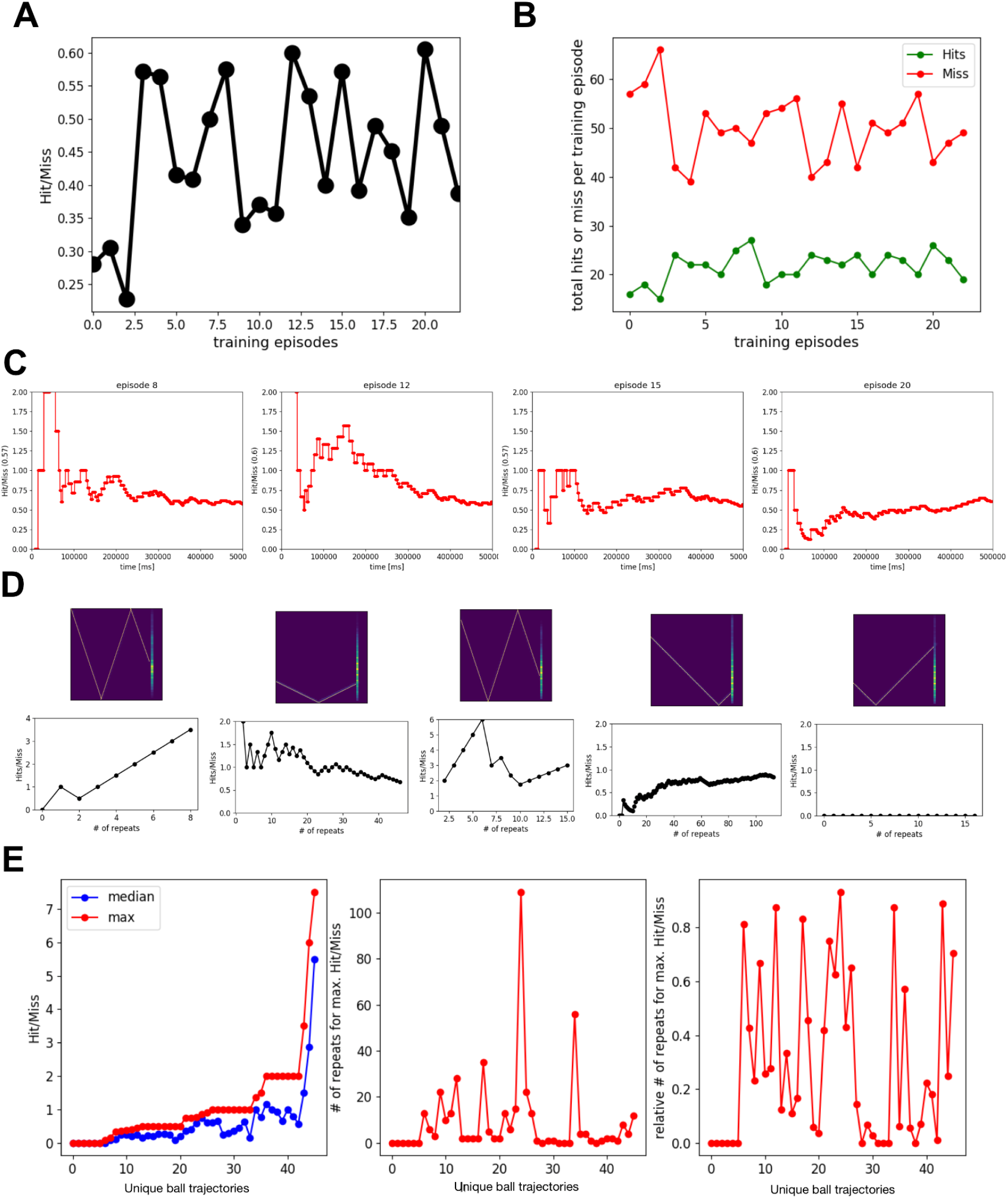
Performance of the recurrent model with *retrograde targeted RL*. **A)** cumulative performance of 23 training episodes. **B)** Cumulative Hits and Misses of 40 training episodes. **C)** Temporal evolution of performance during training episodes 8, 12, 15, 20 (selected arbitrarily). **D)** Examples of ball trajectories with model’s learning performance shown for different modes of learning. First example shows a ball trajectory for which the model kept learning. Second example shows a ball trajectory for which the model performed well at the beginning of the training and then kept forgetting as shown by decrease in hit to miss ratio. In the third example, the model learned in the beginning and then forgot and then again started learning. Fourth example showed sustained performance as the model’s ability to hit the ball plateaued and remained constant for 80 repeats. In the fifth example, the model was unable to learn how to hit the ball for this ball trajectory. **E)** The model’s performance for different ball trajectories: The left panel shows the median and maximum Hit/Miss values during repeated occurrences of the unique ball trajectories. The middle panel shows the number of repeats of the unique ball trajectories at which the model showed peak performance. The right panel shows the relative number of repeats of the unique ball trajectories at which the model showed the peak performance. This indicates that for some ball trajectories (# 30-32), the model performed best without any training and the training only reduced the performance of the model. For some ball trajectories (seq # 0-5), the model could not learn to hit the ball. This also shows that for some ball trajectories (see the seqs with relative # of repeats for max. Hit/Miss values between 0.2 and 0.8), the model first learns to hit the ball and then forgets, whereas for a few ball trajectories (see the seqs with relative # of repeats for max. Hit/Miss values 0.8 or above), the model did not forget how to hit the ball until the end of all training episodes.

**Supplementary Figure 3.**
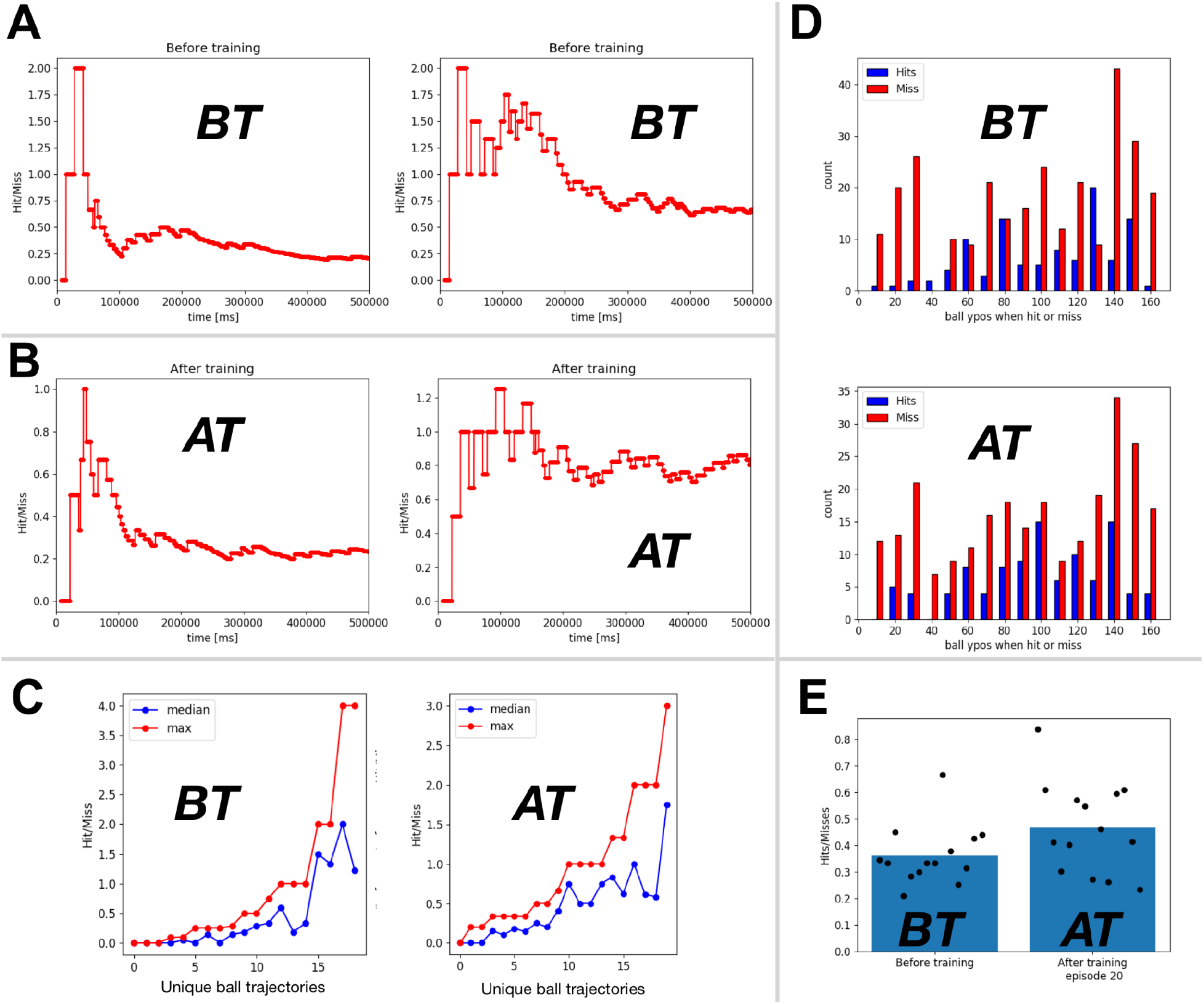
The recurrent spiking neuronal network model using retrograde targeted RL showed variable performance after learning. A) Temporal evolution of the performance of two example simulations using different initial positions of the racket and the ball and initial weights for synaptic connections. B) same as in A using weights after training episode 20 for synaptic connections. C) Summary of the peak and the median performance for all different ball trajectories for the model before training and using weights after training episode 20. D). The histogram of ‘Hits’ and ‘Misses’ against the ball’s vertical position (ypos) when crossing the racket for the model before and after training E) The bar plot shows the mean (n=14; filled circles) performance (Hit/Miss) of the model before training (using initial weights), after training episode 20. For the performance comparison, we ran 500 sec simulations using 14 different initial positions of the racket and the ball.

